# A network correspondence toolbox for quantitative evaluation of novel neuroimaging results

**DOI:** 10.1101/2024.06.17.599426

**Authors:** Ru Kong, R. Nathan Spreng, Aihuiping Xue, Richard F. Betzel, Jessica R. Cohen, Jessica S. Damoiseaux, Felipe De Brigard, Simon B. Eickhoff, Alex Fornito, Caterina Gratton, Evan M. Gordon, Avram J. Holmes, Angela R. Laird, Linda Larson-Prior, Lisa D. Nickerson, Ana Luísa Pinho, Adeel Razi, Sepideh Sadaghiani, James M. Shine, Anastasia Yendiki, B. T. Thomas Yeo, Lucina Q. Uddin

**Affiliations:** Centre for Translational MR Research and Centre for Sleep & Cognition, National University of Singapore, Singapore; Department of Neurology and Neurosurgery, McGill University, Montreal, Quebec, Canada; Department of Psychological and Brain Sciences, Indiana University, Bloomington, IN, USA; Department of Psychology and Neuroscience, University of North Carolina, Chapel Hill, NC, USA; Institute of Gerontology and Department of Psychology, Wayne State University, Detroit, MI, USA; Department of Philosophy, Duke University, Durham, NC, USA; Institute of Systems Neuroscience, Heinrich Heine University Düsseldorf, Düsseldorf, Germany; School of Psychological Sciences, Turner Institute for Brain and Mental Health, and Monash Biomedical Imaging, Monash University, Melbourne, Victoria, Australia; Department of Psychology, Florida State University, Tallahassee, FL, USA; Mallinckrodt Institute of Radiology, Washington University, St. Louis, MO, USA; Department of Psychiatry, Center for Brain Health, Rutgers University, New Brunswick, NJ, USA; Department of Physics, Florida International University, Miami, FL, USA; Department of Psychiatry and Department of Neurobiology and Developmental Sciences, University of Arkansas for Medical Sciences, Little Rock, AR, USA; Department of Psychiatry, Harvard Medical School, McLean Hospital, Boston, MA, USA; Brain and Mind Institute, Western University, London, Ontario, Canada; Department of Psychology and Beckman Institute for Advanced Science and Technology, University of Illinois, Urbana Champaign, IL, USA; Brain and Mind Center, University of Sydney, Sydney, New South Wales, Australia; Department of Radiology, Massachusetts General Hospital, Boston, MA, USA; Department of Psychiatry and Biobehavioral Sciences and Department of Psychology, University of California Los Angeles, CA, USA

## Abstract

Decades of neuroscience research has shown that macroscale brain dynamics can be reliably decomposed into a subset of large-scale functional networks, but the specific spatial topographies of these networks and the names used to describe them can vary across studies. Such discordance has hampered interpretation and convergence of research findings across the field. To address this problem, we have developed the *Network Correspondence Toolbox* (NCT) to permit researchers to examine and report spatial correspondence between their novel neuroimaging results and sixteen widely used functional brain atlases, consistent with recommended reporting standards developed by the Organization for Human Brain Mapping.

The atlases included in the toolbox show some topographical convergence for specific networks, such as those labeled as “default” or “visual”. Network naming varies across atlases, particularly for networks spanning frontoparietal association cortices. For this reason, quantitative comparison with multiple atlases is recommended to benchmark novel neuroimaging findings.

We provide several exemplar demonstrations using the Human Connectome Project task fMRI results and UK Biobank independent component analysis maps to illustrate how researchers can use the NCT to report their own findings through quantitative evaluation against multiple published atlases. The NCT provides a convenient means for computing Dice coefficients with spin test permutations to determine the magnitude and statistical significance of correspondence among user-defined maps and existing atlas labels. The NCT also includes functionality to incorporate additional atlases in the future. The adoption of the NCT will make it easier for network neuroscience researchers to report their findings in a standardized manner, thus aiding reproducibility and facilitating comparisons between studies to produce interdisciplinary insights.

## Introduction

Standardized scientific nomenclature can facilitate the effective communication of concepts related to complex biological systems. Neuroscience has a long history of classifying distinct anatomical regions of the brain across multiple spatial scales, from the cortical lobes, discrete gyri and sulci of the cerebral cortex and subcortical nuclei, to Brodmann’s cytoarchitectonic areas ^1^. Attempts have also been made to categorize well-defined neural circuits and systems based on various anatomical and functional criteria, such as the cortico- spinal tract, cortico-striato-thalamic circuits, and the dorsal and ventral visual processing streams^2^. Such attempts at standardization facilitate communication about brain organization between researchers and assist in educating students.

Modern neuroimaging methods, particularly those relying on magnetic resonance imaging (MRI), have revolutionized our capacity to non-invasively map different aspects of brain structure and function in living humans. A key development has been the ability to map functional brain networks, or groups of brain areas that exhibit functional synchrony. As a result of the growing popularity of this approach, many findings from neuroimaging studies are described in terms of the *brain networks,* rather than the *brain areas,* where they are localized. However, this is often done in an ad hoc way, due to the lack of standardization in the naming of these networks. This inconsistency in scientific reporting complicates comparisons of findings across studies and limits the integration of novel discoveries ^3^.

The emergence of functional brain network atlases, which are now routinely used in cognitive and network neuroscience ^4^, should, in principle, contribute to standardization in the naming of functional networks. These atlases typically assign parcellated cortical and/or subcortical regions to one of a set of large-scale brain networks with varying topographies, ranging from largely contiguous unimodal territories (e.g. visual network in occipital cortex) to associative systems that span multiple spatially segregated and distributed regions across the brain (e.g., default network encompassing frontal, temporal and parietal regions). The atlases also vary in how they name large-scale brain networks and in terms of the number of brain networks, as well as their approach to defining the networks, characterizing their spatial topography, and deciding how they should be labeled (e.g., based on their anatomical location or purported cognitive correlates) ^5^. These differences between atlases lead to further variability and uncertainty in scientific reporting.

Recognizing these challenges, the Organization for Human Brain Mapping (OHBM) established a *Best Practices* committee on large-scale brain network nomenclature, following the success of earlier similar consensus-building initiatives aiming to enhance standardization by providing reporting guidelines for the field ^6,7^. This committee, formed in 2020, is now known as the Workgroup for HArmonized Taxonomy of NETworks (WHATNET). Initial attempts by the workgroup to conceptually identify convergence across large-scale functional brain networks led to discussion and enumeration of the multiple challenges inherent to this enterprise ^8^. The workgroup surveyed hundreds of scientists and solicited putative network names based upon renderings of various network spatial topographies. Results from the survey, described in detail in the initial report from WHATNET ^8^ and publicly available (https://osf.io/3fzta/), indicated that networks based in non-distributed unimodal brain regions (somatomotor and visual systems) could be readily and reliably identified. The somatomotor network showed 97% agreement amongst raters, while the visual network showed 92% agreement amongst raters. Only one spatially distributed network, the default network, demonstrated consensus agreement in name and topography, with 93% agreement amongst raters. Other networks, such as those sometimes referred to as salience or frontoparietal, showed less consistency. These findings indicate that consensus agreement in nomenclature is limited to the extreme poles of the canonical cortical hierarchy ^9^, anchored at one end by the visual and somatomotor systems and on the other by the default network. In between these systems is an expanse of cortex with observable network structure, but with little agreement between researchers regarding their spatial topography or nomenclature. Such variability poses a major problem for any attempts to compare or integrate findings across studies.

In response to the WHATNET survey findings, here we present the *Network Correspondence Toolbox* (NCT). This toolbox provides researchers with a practical solution for labeling novel observed fMRI activation, functional connectivity patterns, or other thresholded neuroimaging maps based on a quantitative evaluation against currently published atlases. The NCT also allows users to assess the spatial correspondence between networks in new and existing atlases. We demonstrate the utility of the NCT in providing network labels to exemplar neuroimaging maps based on a quantitative evaluation of spatial correspondence with multiple published network atlases. Quantitative evidence of correspondence is determined by the magnitude of the Dice coefficient, in addition to spin test permutations for robust statistical assessment of significance. We conclude with a recommendation that future network and cognitive neuroscience reports include an evaluation of the correspondence between their network labeling schemes and multiple published network atlases produced by independent research laboratories, as facilitated by the NCT. This approach transparently acknowledges and quantitatively addresses the ambiguity inherent in assigning labels to topographic brain maps and encourages greater alignment across network neuroscience studies by objectively assessing the convergence or divergence between new findings and published network labels and schema.

Importantly, the NCT is not prescriptive regarding the largely intractable problem of network nomenclature. Rather, it is a tool to facilitate comparison and interpretation to be drawn across atlases, ultimately resulting in greater standardization in reporting of results.

## Results

### Network Correspondence Toolbox

Anatomical localization of patterns of brain activity is tedious ^10^, but many digital and paper atlases are available to facilitate the process ^11^.

Localization to large-scale functional brain networks is a more recent development, with standardized tools in demand (see ^12^ for a recent introduction of probabilistic network localization). The NCT flexibly permits the assessment of spatial correspondence between several target reference atlases and novel brain maps including, but not limited to, functional connectivity and task activation results, as well as other maps in standard space ^13^. The first release of the NCT includes sixteen widely used functional brain atlases; note that some of the included atlases contain multiple versions: Yeo2011, Schaefer2018, Kong2021, Power2011, Shen2013, Gordon2017, Cole-Anticevic2019, ICA-UK Biobank, ICA-HCP, ICA-BrainMap and Shirer2012 ^14–25^ (**Figure 1, Supplementary Table 1**).

**Figure 1.**
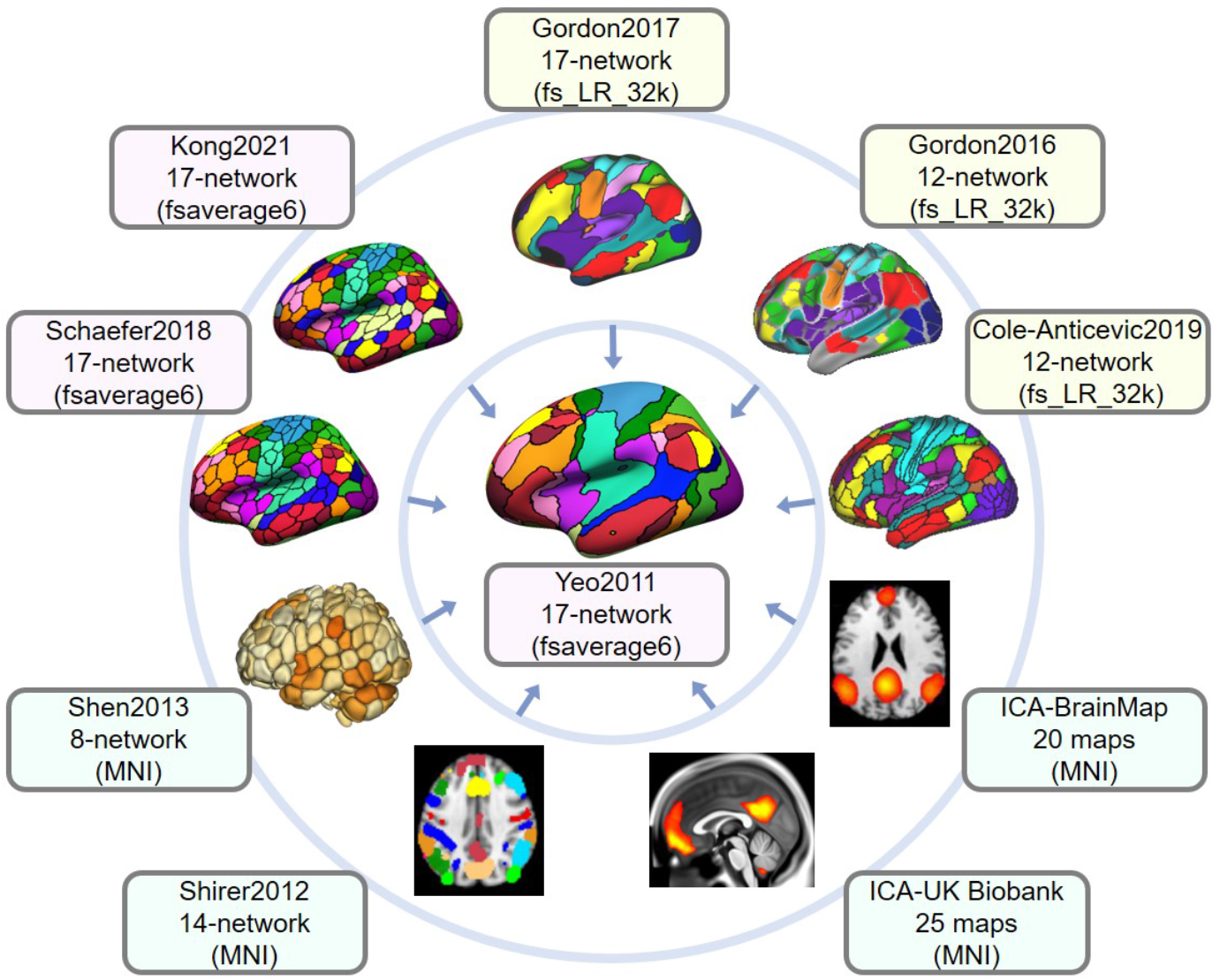
Example atlases included in the Network Correspondence Toolbox (NCT). The NCT is a toolbox that facilitates exploration of network correspondences across multiple functional network atlases as well as quantitative comparison of novel neuroimaging results with multiple atlases. Ten atlases are shown here for illustration purposes. In this example, the Yeo 17-network atlas in fsaverage6 space (center) serves as the reference atlas. All other surrounding atlases in different spaces are projected to the fsaverage6 space to compute Dice overlap coefficients with the reference networks. A full list of available atlases in the NCT can be found in **Supplementary Table 1**.

First, we provide step-by-step guidelines for NCT usage. Next, as a representative example, we demonstrate the correspondence between two widely-used atlases that delineate the cortex into 17 discrete networks to motivate NCT usage: Yeo2011-17 and Gordon2017-17.

Finally, we report several examples of how the NCT can be used by treating publicly available Human Connectome Project (HCP) task fMRI results and UK Biobank independent component analysis (ICA) maps as “novel” findings for demonstration purposes. These specific maps were chosen to illustrate how the NCT performs in cases where the input data correspond with existing networks to a strong degree, as well as in more challenging scenarios where correspondence is lower.

### How to use the NCT

For each network of a reference atlas, the magnitude of spatial correspondence with an empirical brain map, and the statistical significance of this correspondence, can be determined using the NCT. All atlases and fMRI results are projected from a standard space into fsaverage6 space. Correspondence is assessed as the Dice coefficient, where 0 is no spatial correspondence and 1 is total spatial correspondence. In order to assess statistical significance, a spin test ^20,26,27^ implemented in Python ^28^ is performed with 1000 permutations for each reference atlas network.

The NCT can be installed using the command *pip install cbig_network_correspondence*. **Figure 2** illustrates the usage of the NCT to explore network correspondence between user-defined input data and a set of atlases. Specifically, users specify the pathway to the input data and provide a configuration file containing input data details such as the data name, data space, and data type. Additionally, users provide a list of atlases included in the NCT, available at https://github.com/rubykong/cbig_network_correspondence#atlases-we-included.

**Figure 2.**
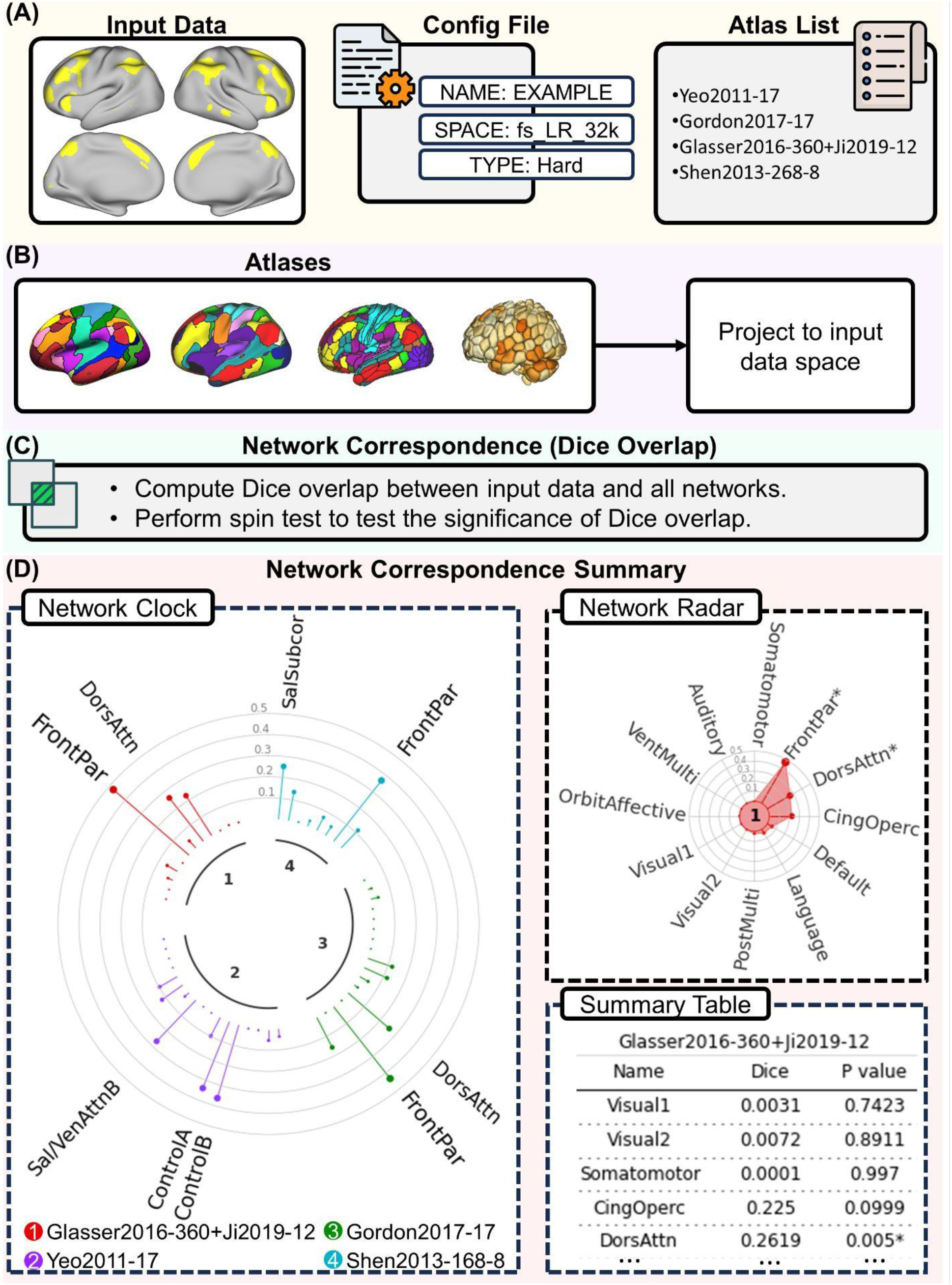
Illustration of NCT usage to explore network correspondence between user-defined input data and a set of atlases. In this example, we explore the overlap between a single-dimension input dataset and 4 atlases: the Yeo2011 17-network atlas (“Yeo2011-17”, the Gordon2017 17-network atlas (“Gordon2011-17”), the Glasser2016 360-ROI atlas with Ji2019 12 Cole-Anticevic networks (“Glasser2016-360+Ji2019-12”), and the Shen2013 268-ROI atlas with 8 networks (“Shen2013-268-8”). (A) The user provides the input data together with a config file specifying the name, data space (e.g., fs_LR_32k, fsaverage6, FSLMNI2mm), and data type (“Metric” if the input data contains floating values; “Hard” if the input data contains binary values). In this example, the input data is in fs_LR_32k space and contains binary values. The data type is “Hard”. The user also provides an atlas list indicating which atlases to include. (B) The NCT reads the input data and projects the atlases in the atlas list to the input data space. (C) The NCT computes the Dice overlap between the input data and networks from the atlases in the list. The NCT also performs spin tests to test whether the input data and networks significantly overlap. (D) The NCT reports the network correspondence summary using a “Network Clock” plot, a “Network Radar” plot, and a summary table for single-dimensional data. The “Network Clock” provides a visual comparison of the Dice overlap across networks from different atlases. Different colors represent different atlases. The lollipop bars represent the Dice overlap coefficients. Networks significantly overlapping with the input data (*p* < 0.05) are indicated by the network names. A larger font size represents a larger Dice coefficient. The “Network Radar” shows the Dice overlap across networks within each atlas. Networks significantly overlapping with the input data (*p* < 0.05) are indicated by “*”. The NCT also provides a summary table showing the exact Dice coefficients and *p*-values across different atlases. The NCT uses network names from the original paper for each atlas.

The NCT proceeds by first reading the input data, projecting the atlases specified in the list to the input data space, computing the spatial similarity between the data and atlases, and conducting statistical analyses. The network correspondence results are then visually presented.

The input data can be either uni-dimensional or multidimensional. Examples of unidimensional data include a spatial map of a single ICA component, a task contrast map, or a single network from a hard parcellation. The network correspondence results of unidimensional data are illustrated as a “Network Clock” plot, a “Network Radar” plot, and a summary table (**Figure 2D**). The “Network Clock” provides a visual comparison of the Dice overlap between the input data and networks from different atlases. Different colors represent different atlases. The lollipop bars represent the Dice overlap coefficients. Networks significantly overlapping with the input data (*p* < 0.05) are indicated by the network names. In this plot, networks that have larger Dice coefficients are shown by larger font sizes. The “Network Radar” shows the Dice overlap across networks within each atlas. Networks significantly overlapping with the input data (*p* < 0.05) are indicated by “*”. The NCT provides a “Summary Table” showing the exact Dice coefficients and *p*-values for different atlases.

In the case of multidimensionality, examples could include a set of ICA spatial maps, multiple task contrast maps, or any atlas included in the NCT. The NCT reports the network correspondence results of multidimensional data as “Overlap Heatmap” plots (**Figure 3D**), where the rows correspond to the input data, and columns correspond to the networks of a comparison atlas. The *k*-th row and *m*-th column represents the Dice overlap coefficient between the *k*-th dimension of the input data and the network *m* from the atlas. In this plot, a brighter color indicates higher overlap, while a darker color indicates lower overlap. Networks significantly overlapping with the input data (*p* < 0.05) are indicated by “*”. The NCT also provides a summary table showing the exact Dice coefficients and *p*-values.

**Figure 3.**
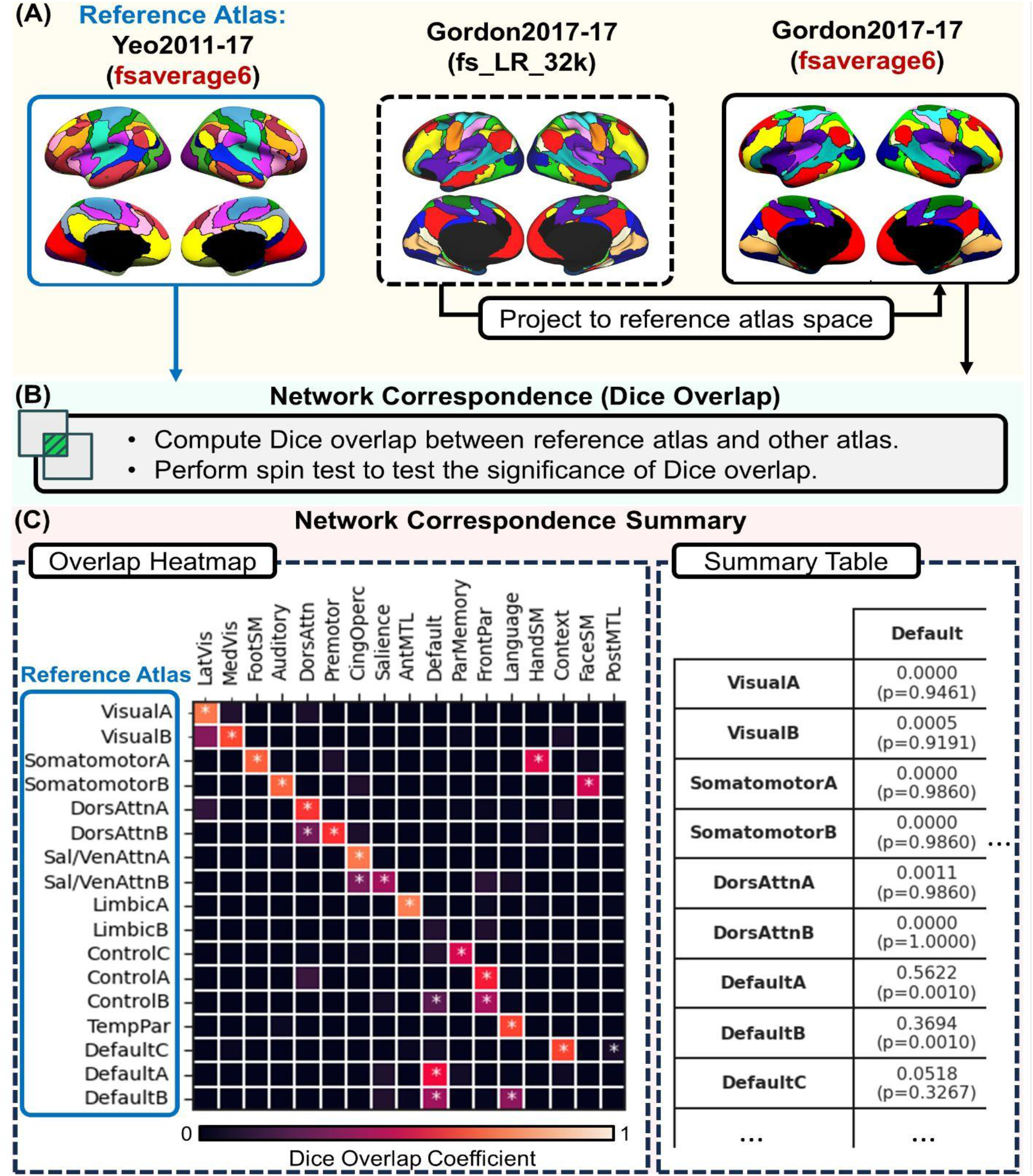
Illustration of NCT usage to explore network correspondence between two atlases. In this example, we explore overlap between the Yeo2011 17-network atlas (reference atlas) and the Gordon2017 17-network atlas. (A) The user specifies the name of the reference atlas (here, Yeo2011-17) and the name of comparison atlas (here, Gordon2017-17). The Yeo 17- network atlas is in fsaverage6 space, while the Gordon2017 17-network atlas is in fs_LR_32k. In this case, the reference atlas space is fsaverage6. Therefore, the NCT projects the Gordon2017 17-network to fsaverage6 space. (B) The NCT computes the Dice overlap between networks from Yeo 17-network atlas and Gordon2017 17-network atlas. The NCT also performs a spin test to test whether networks significantly overlap. (C) The NCT reports the network correspondence between these two atlases using an overlap heatmap plot, where the *k*-th row and *m*-th column represents the Dice overlap coefficient between network *k* of the Yeo 17-network atlas and network *m* from the Gordon2017 17-network atlas. A high Dice coefficient indicates high overlap between the two networks. Brighter colors indicate higher overlap, darker colors indicate lower overlap. The “*” indicates two networks significantly overlap (*p* < 0.05). Most Yeo2011 networks overlap with at least one network in the Gordon2017 atlas. The NCT also provides a summary table showing the exact Dice coefficients and *p-*values (see **Supplementary Table 2**). The NCT uses network names from the original paper for each atlas.

The NCT was built in Python and is available at the GitHub repository: https://github.com/rubykong/cbig_network_correspondence. The toolbox has been released to the Python Package Index (PyPI): https://pypi.org/project/cbig-network-correspondence.

Detailed documentation and a tutorial are available at: https://rubykong.github.io/cbig_network_correspondence. *Example Atlas Network Correspondence* As a quantitative demonstration of the network correspondence problem that motivated the development of the NCT, we evaluated network-to-network spatial correspondence between two widely used atlases. For this illustration, all 136 network pairs from the Yeo2011-17 and Gordon2017-17 atlases were examined (**Figure 3**; See **Supplementary Table 2** for Dice coefficients and corresponding *p*-values). Overall, we observed good correspondence between several of the networks across these two atlases. For example, Gordon2017-17 Lateral Visual and Medial Visual networks significantly corresponded to the Yeo2011-17 Visual A and B networks, respectively. Specific Hand, Face, and Foot Somatomotor networks in the Gordon2017-17 showed significant correspondence with Somatomotor networks A and B in Yeo2011-17. However, notable discrepancies also emerged, particularly with regard to network name and purported function. For example, the Gordon2017-17 Auditory network also spatially corresponded to the Yeo2011-17 Somatomotor network B. While the Gordon2017-17 Dorsal Attention network showed significant correspondence with the Yeo2011-17 Dorsal Attention networks A and B, the Gordon2017-17 Premotor network also significantly corresponded with the Yeo2011-17 Dorsal Attention network B. The Gordon2017-17 Cingulo-opercular network significantly corresponded with the Yeo2011-17 Salience / Ventral Attention networks A and B. Yet a distinct Gordon2017-17 Salience network also corresponded with Yeo2011-17 Salience / Ventral Attention network B. The Gordon2017-17 Anterior Medial Temporal Lobe network significantly corresponded to Yeo2011-17 Limbic A network (which comprises the anterior temporal lobes). In robust agreement, the Gordon2017-17 Frontoparietal network significantly corresponded to the Yeo2011-17 Frontoparietal Control networks A and B. However, the Gordon2017-17 Parietal Memory network significantly corresponded to the Yeo2011-17 Frontoparietal Control network C. In another example of seemingly good spatial alignment, the Gordon2017-17 Default network significantly corresponded to the Yeo2011-17 Default networks A and B. However, the Gordon2017-17 Default network also showed significant correspondence with Yeo2011-17 Frontoparietal Control network B, highlighting the importance of considering the relative magnitude of the Dice coefficient. The Gordon2017-17 Language network significantly corresponded with the Yeo2011-17 Temporal Parietal network, as well as Default network B. The Gordon2017-17 Context network and Posterior Medial Temporal Lobe network both significantly corresponded to the Yeo2011-17 Default network C. Notably, the Yeo2011-17 Limbic B network (which comprises orbitofrontal cortex) showed no correspondence with any Gordon2017-17 networks.

This Yeo2011-17 and Gordon2017-17 example illustrates the problems of 1) incongruence in network nomenclature, and 2) varying levels of network correspondence across atlases. A pair-wise spatial correspondence analysis of all networks included within all sixteen atlases is outside the scope of the present report. However, the NCT retains this functionality for interested readers. In-depth pairwise assessment between brain network atlases, in addition to benchmarking of novel neuroimaging results against networks defined across multiple atlases, is a core function of the NCT. Due to spatial and nomenclature variability across atlases, the OHBM *Best Practices* committee now recommends that users use multiple atlases originating from different research laboratories for reporting correspondence between novel results and existing network labels. The recommendation to report results in reference to multiple atlases comes from the observation that even networks with similar nomenclature across atlases (e.g. “salience”) may not share the same spatial topography (**Figure 4**).

**Figure 4.**
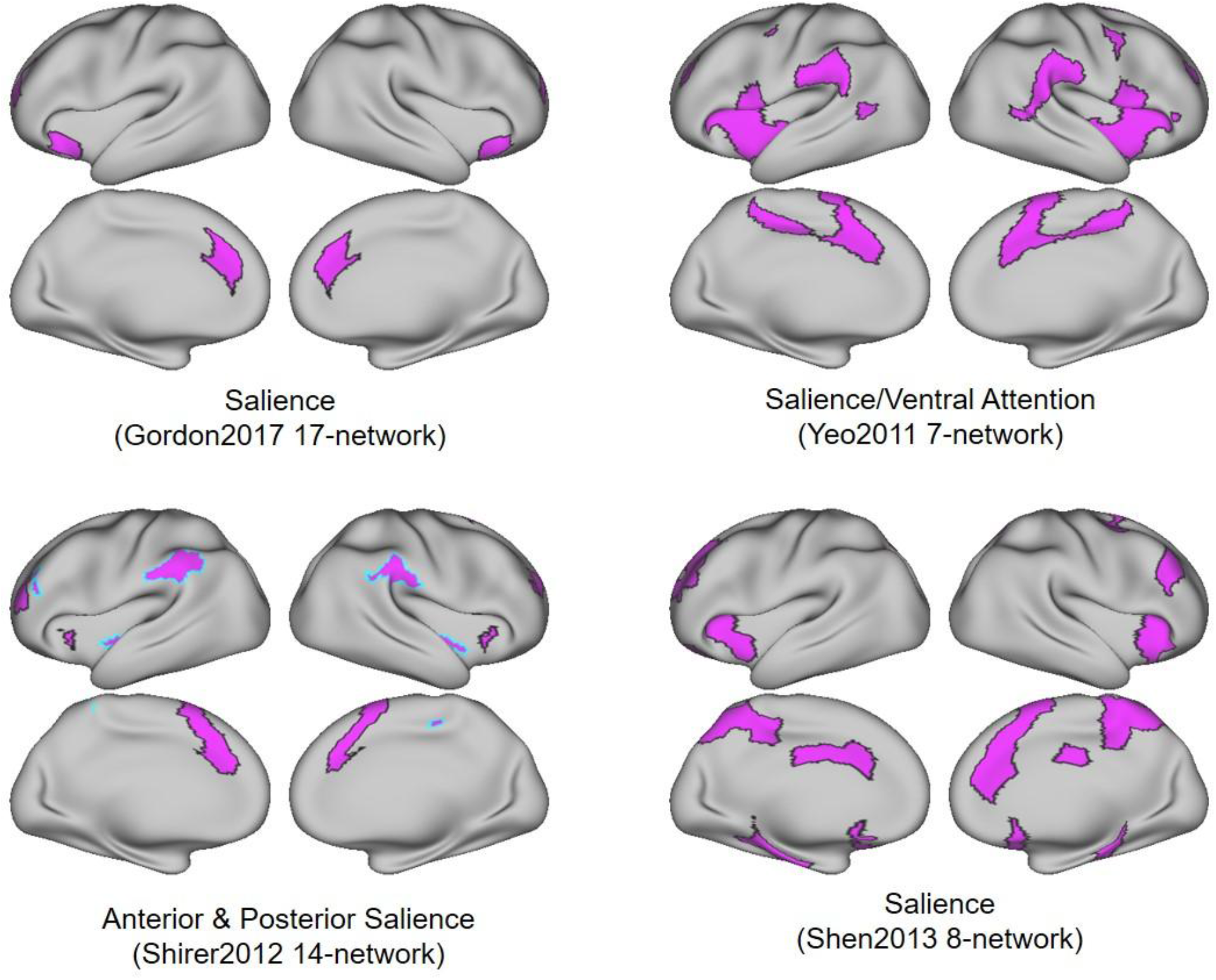
Networks with similar names can show different spatial topographies. “Salience” networks from four different atlases. These networks are labeled using similar nomenclature across multiple atlases, though they span different, largely non-overlapping anatomical locations.

### Illustrating the utility of the NCT

To demonstrate various applications of the NCT in both straightforward and challenging cases, we next show multiple network correspondence results. For the following examples, the image from an analysis (e.g., task activation map) was loaded, the space of the image set to a specific standard space (e.g., MNI152), and a threshold set for significant voxels. Dice coefficients were then determined for all reference atlases, and output as a Dice coefficient array. For each network within each atlas, we determined the Dice coefficient of network overlap and used spin tests to quantify levels of significance.

Networks significantly overlapping with the thresholded input data (*p* < 0.05) are indicated by the network names. Larger font sizes represent larger Dice coefficients. The “Network Radar” shows the Dice overlap across networks within each atlas. Networks significantly overlapping with the thresholded input data (*p* < 0.05) are indicated by “*” in the “Network Radar”.

For the first example, we used the HCP working memory task fMRI contrast (2BK-0BK) which includes data from 997 HCP S1200 subjects ^29^. As illustrated in the “Network Clock” depicted in **Figure 5**, this task fMRI contrast (top left) shows significant spatial overlap with several networks across multiple atlases (e.g. “FrontPar” network from the Glasser and Gordon atlases, “DorsAttn” network from the Laumann atlas). As shown in the “Network Radar” plot (top right), the NCT can be further used to quantify the overlap between user-defined maps and specific networks within an atlas. In the example highlighted in this figure, the input contrast shows significant overlap with the network labels “FrontPar” and “DorsAttn” in the Gordon atlas. This example illustrates a case where the task activation map corresponds well with networks specified in multiple atlases.

**Figure 5.**
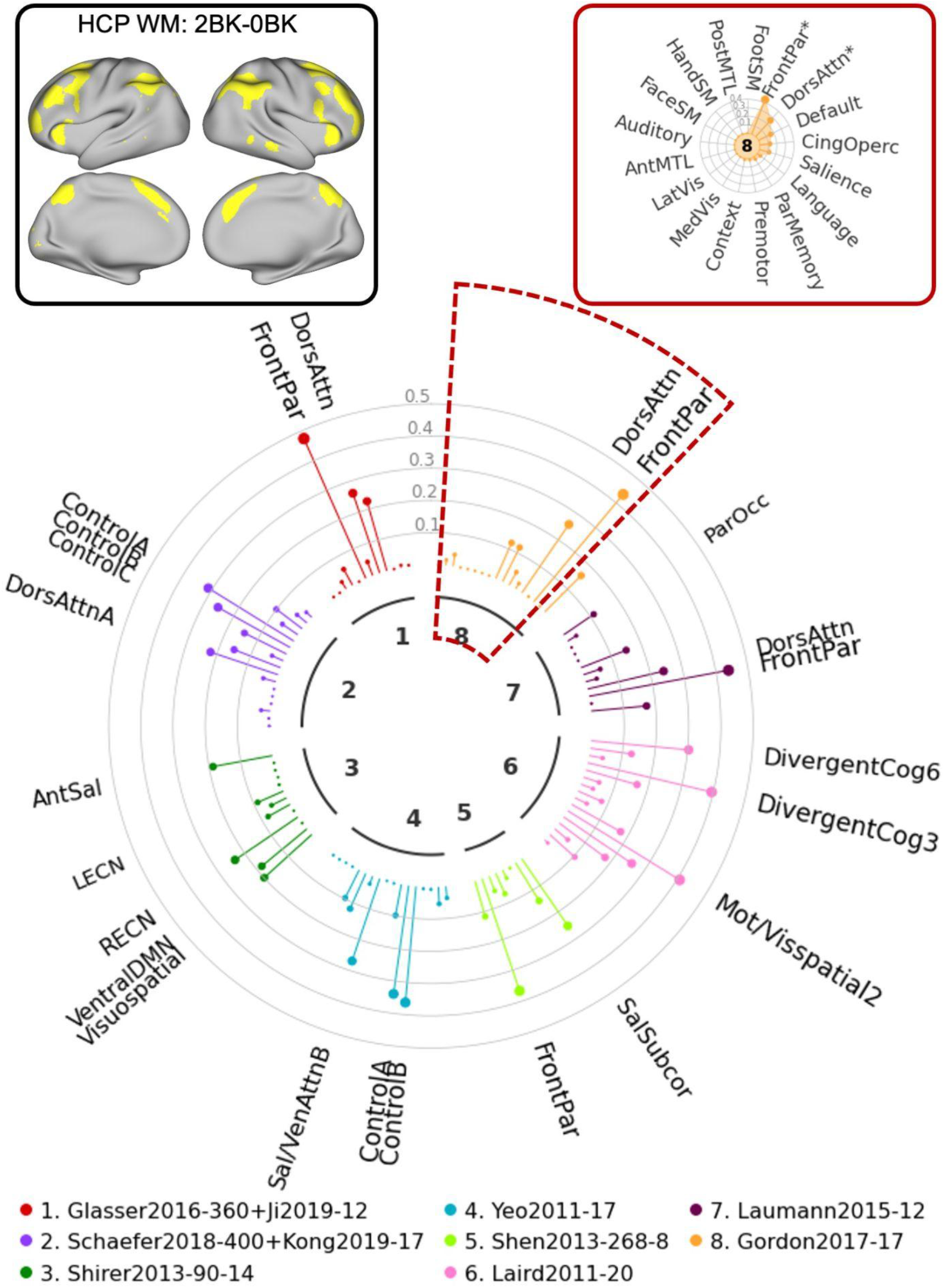
Illustration of usage of the NCT to explore network correspondence between user-defined input data and multiple atlases: HCP Example 1. In this example, we explore overlap between the HCP working memory task contrast (2BK vs. 0BK) and networks from 8 atlases. The user provides the input data together with a config file specifying the name, data space, data type, and the threshold which is used to threshold the input data. The user also provides an atlas list indicating which atlases to include. The NCT thresholds the data based on the config file and projects the atlases in the atlas list to the input data space. The NCT computes the Dice overlap between the thresholded data and all networks from atlases in the list. The NCT also performs spin tests to test whether the thresholded input data and networks significantly overlap. The NCT reports the network correspondence summary using a “Network Clock” plot (middle), a “Network Radar” plot (top right), and a summary table showing the exact Dice coefficients and *p*-values for all atlases (see **Supplementary Table 3**). The “Network Clock” provides a visual comparison of Dice overlap across networks from different atlases. Different colors represent different atlases. The lollipop bars represent the Dice overlap coefficients.

For the second example, we used the HCP social task fMRI contrast (TOM-random) which includes data from 997 subjects. As illustrated in the “Network Clock” depicted in **Figure 6**, this task fMRI contrast (top left) shows significant spatial overlap with several networks across multiple atlases (e.g. “VentAttn” network from the Laumann atlas, “Language” network from the Gordon atlas). As shown in the “Network Radar” plot (top right), The NCT further can be used to quantify the overlap between user-defined maps and specific networks within an atlas. In the example highlighted in this figure, the input contrast shows significant overlap with the network labels “Language” and “Context” in the Gordon atlas. This example illustrates a case where the task activation map does not correspond well with networks specified in multiple atlases. It thus signals that any network labels should be applied to this map with great caution.

**Figure 6.**
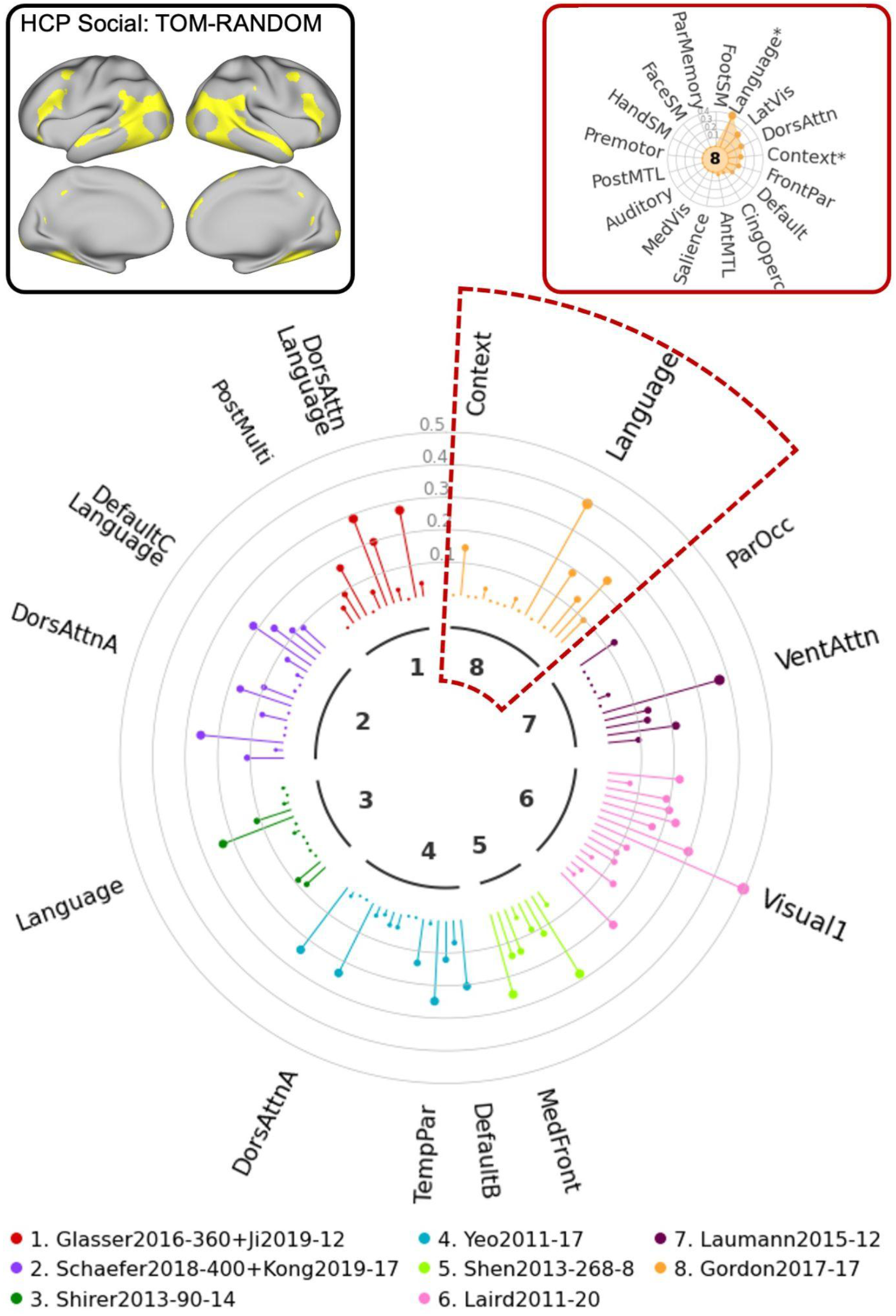
Illustration of NCT usage to explore network correspondence between user-defined input data and multiple atlases: HCP Example 2. In this example, we explore the overlap between the HCP social task contrast (theory of mind vs. random) and 8 atlases. The user provides the input data together with a config file specifying the name, data space, data type, and the threshold which is used to threshold the input data. The user also provides an atlas list indicating which atlases to include. The NCT thresholds the data based on the config file and projects the atlases in the atlas list to the input data space. The NCT computes the Dice overlap between the thresholded data and all networks from atlases in the list. The NCT also performs spin tests to test whether the thresholded input data and networks significantly overlap. The NCT reports the network correspondence summary using a “Network Clock” plot (middle), a “Network Radar” plot (top right), and a summary table showing the exact Dice coefficients and *p*-values for all atlases (see **Supplementary Table 4**). The “Network Clock” provides a visual comparison of Dice overlap across networks from different atlases. Different colors represent different atlases. The lollipop bars represent the Dice overlap coefficients. Networks significantly overlapping with the thresholded input data (*p* < 0.05) are indicated by the network names. Larger font sizes represent larger Dice coefficients. The “Network Radar” shows the Dice overlap across networks within each atlas. Networks significantly overlapping with the thresholded input data (*p* < 0.05) are indicated by “*” in the “Network Radar”.

For the next example, we used the UK Biobank ICA component 5 to illustrate how the NCT can facilitate reporting of results from this type of analysis. As illustrated in the “Network Clock” depicted in **Figure 7**, this ICA map (top left) shows significant spatial overlap with networks labeled “Visual” across multiple atlases. The final example (**Figure 8**) uses UK Biobank ICA component 3 as the input map. Even with this complex input spanning multiple brain areas, the NCT can be used to quantify overlap with existing network labels to facilitate communication and comparison. Again, these two examples illustrate cases in which the input maps correspond well (**Figure 7**) and do not correspond well (**Figure 8**) with networks specified in multiple atlases.

**Figure 7.**
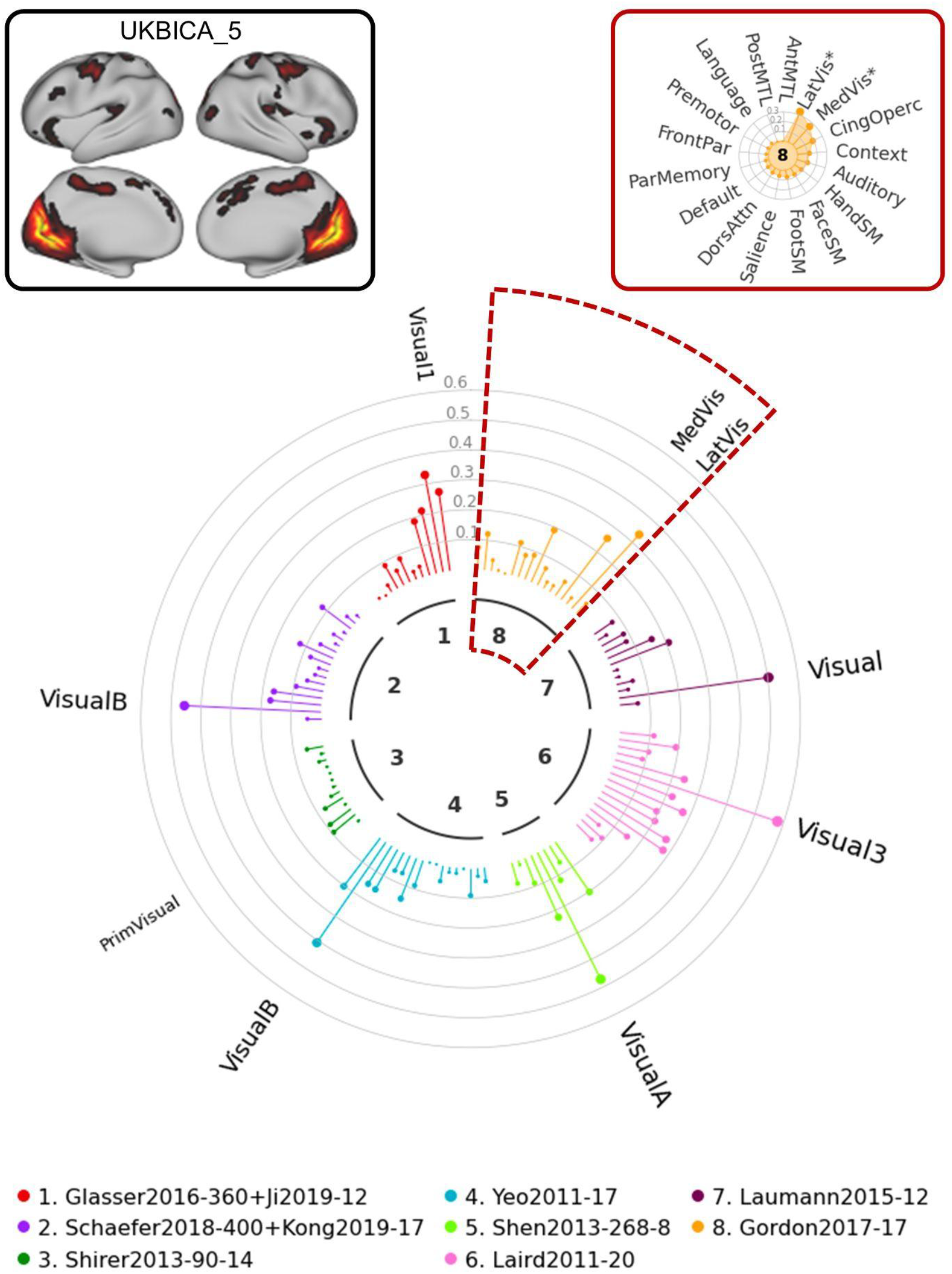
Illustration of NCT usage to explore network correspondence between user-defined input data and multiple atlases: UKB Example 1. In this example, we explore overlap between UKB ICA component 5 and 8 representative atlases. The UKB ICA z-stat maps with 21 good components (https://www.fmrib.ox.ac.uk/ukbiobank/group_means/rfMRI_GoodComponents_d25_v1.txt) were thresholded by FSL melodic mixture-modeling threshold 0.6. The component 16 corresponds to the cerebellum and was further excluded, resulting in 20 thresholded UKB ICA maps. The 20 thresholded UKB ICA maps can be found in **Supplementary** Figure 1. The NCT also provides a summary table showing the exact Dice coefficients and *p*-values for all atlases (**Supplementary Table 5**).

**Figure 8.**
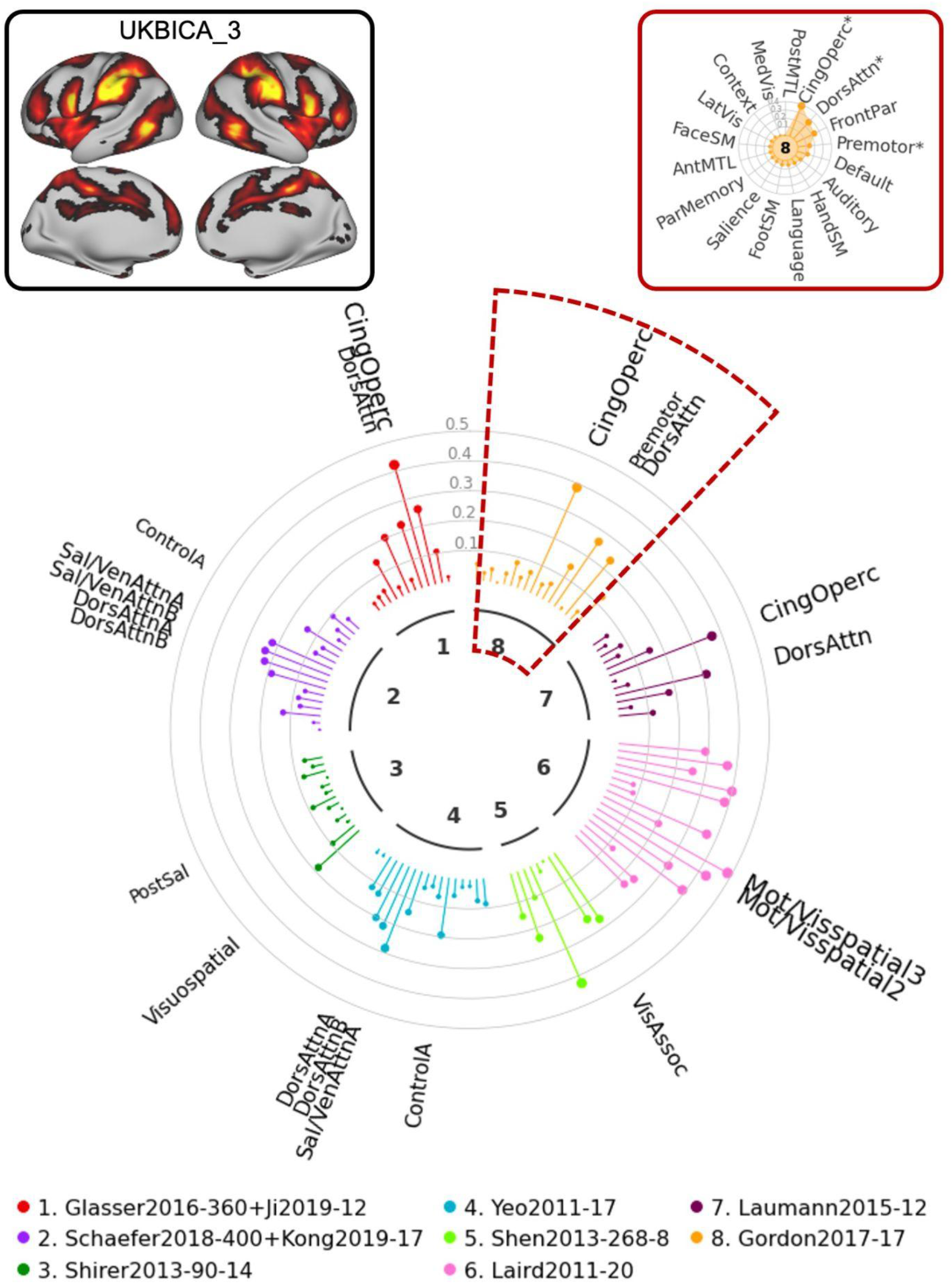
Illustration of NCT usage to explore network correspondence between user-defined input data and multiple atlases: UKB Example 2. In this example, we explore overlap between UKB ICA component 3 and 8 representative atlases. The UKB ICA z-stat maps with 21 good components (https://www.fmrib.ox.ac.uk/ukbiobank/group_means/rfMRI_GoodComponents_d25_v1.txt) were thresholded by FSL melodic mixture-modeling threshold 0.6. The component 16 corresponds to the cerebellum and was further excluded, resulting in 20 thresholded UKB ICA maps. The 20 thresholded UKB ICA maps can be found in **Supplementary** Figure 1. The NCT also provides a summary table showing the exact Dice coefficients and *p*-values for all atlases (**Supplementary Table 6**).

Taken together, these examples illustrate how researchers can use the NCT to quantitatively and accurately present novel cognitive and network neuroscience results in reference to widely used atlases to enhance interpretability and comparison with existing findings in the literature.

## Discussion

Despite decades of functional neuroimaging research resulting in repeated and robust observation of multiple large-scale functional brain networks, the nomenclature describing these networks in humans is not standardized ^5^. Instead, there has been a proliferation of different population-average network atlases that rely on distinct nomenclatures derived by independent research groups using disparate approaches ^8^. Indeed, we have seen that even networks with similar nomenclature (e.g., “salience”) across atlases may not share the same spatial topography.

Although atlases vary both in their number of networks and the methods used to identify networks, they all seek to categorize network affiliations for the cortical and subcortical mantle. The goal of each OHBM *Best Practices* committee is to formalize and endorse best practices across domains of human neuroimaging research to promote science literacy, transparency, and reproducibility. Building on the consensus conclusions from the OHBM *Best Practices* committee on network nomenclature (WHATNET), here we introduce the *Network Correspondence Toolbox* (NCT).

In building the NCT we first investigated the spatial correspondence of discrete large-scale functional brain networks defined in two widely-used atlases (Yeo2011-17 and Gordon2017-17). Overall, spatial correspondence across atlases was observed for many functional networks. However, areas of notable divergence across atlases were also observed, especially for networks spanning association cortices. For example, a network presenting as unitary in one atlas was in some cases broken down into two networks in the other atlas.

Network nomenclature across the two atlases was also inconsistent in many cases. Such discrepancies support the published 2023 recommendations of WHATNET advising that network-based interpretations of novel activation or functional connectivity findings use multiple reference atlases produced by independent research laboratories to corroborate network localization ^8^. The NCT is designed to further promote and facilitate the adoption of this recommendation by providing researchers with a user-friendly tool to readily conduct and report on network correspondences, as demonstrated with multiple examples (**Figures 5-8**). We suggest that future studies reference multiple independent atlases to quantitatively evaluate network localization of activity or functional connectivity, using the NCT ^13,30^. This would support cross-study comparisons for replicability, aid in the interpretation of novel findings, and encourage greater dialogue across research groups and the field of network neuroscience more broadly.

Many novel discoveries in cognitive and network neuroscience are contextualized within the bounds of previously identified network topographies. Methods for assessing network overlap with novel functional connectivity or fMRI task activation maps vary from visual inspection, Euclidean distance between network regions’ center of mass and activation peaks, to direct quantification between an atlas and neuroimage. Variability in image formats between atlases and result images can be a significant obstacle to assessing correspondence directly and/or empirically. Many fMRI task studies are conducted in volume space, yet some atlases are only shared by the creators in surface space ^24,31^, arguing that volume rendering is less precise ^32^. In order to surmount this problem, all 16 atlases included in the NCT were transformed from their original standard space to be interoperable and compatible with all other standard spaces, in the volume or surface. This approach was integral to assessing the correspondence between all atlases. Further, this approach now allows for a direct and empirical examination of novel neuroscience findings against multiple existing widely used atlases simultaneously.

Previous work has attempted to quantify spatial variability among major large-scale brain networks as defined in widely used functional atlases, also noting only modest similarity between atlases. Some have created atlas-based labeling tools ^33^, but none have provided a means for comparing novel results with multiple atlases simultaneously. One project created a Consensual Atlas of REsting-state Network (CAREN) based on Yeo, Gordon, and Doucet atlases^34^. Initial efforts to standardize human brain parcellations have resulted in tools such as *Neuroparc*, which provides a means for comparison among widely used atlases ^35^. Others have compared anatomical, functional connectivity, and random parcellations and concluded that there is no optimal approach for choice of parcellation technique ^36^. Recent parcellation evaluation studies have suggested that multi-modal parcellations combining functional and anatomical metrics may perform worse than those based only on functional data because functionally homogeneous brain regions can span major anatomical landmarks ^37^. Critically, these studies have either simply evaluated the correspondence between different atlases or attempted to derive a single ‘consensus’ atlas. The NCT obviates the need to rely on any single atlas. Instead, it transparently addresses the inherent ambiguity in assigning network labels to empirical brain maps by objectively quantifying the spatial correspondence between a given empirical map and multiple existing, widely used atlas nomenclatures simultaneously.

It should be noted that all neuroimaging-derived spatial maps are sensitive to thresholding choices. The issue of appropriate thresholding of fMRI results is outside the scope of the current work, but several previous discussions of this topic are available, and guidelines regarding thresholding are put forth in early OHBM *Best Practices* reports ^6^. We recommend unthresholded image deposition in an open access repository (e.g. Neurovault https://neurovault.org/) so that researchers can come to their own conclusions regarding overlap between novel results and existing parcellation schemes.

The WHATNET now recommends that new scientific reports on patterns of task activation or functional connectivity, if purported to engage a specific large-scale functional brain network, be indexed to multiple independent atlases for verification. The NCT provides an accessible, easy-to-use tool to follow this recommendation in full. Critically, we see the NCT as an evolving resource, wherein the databases of atlases will be open to the research community. This will enable the NCT, as a tool to quantify network correspondences, to expand as novel parcellation schemes and atlases are introduced. Tools like the NCT will be especially critical for integrating findings across psychiatry and neurology, where the lack of guidance regarding atlas correspondence and network nomenclature can be an impediment to clinical progress ^38^.

## Online Methods

### Functional brain atlases

We investigated cortical network correspondences between user-defined input data and a set of functional network atlases. We considered 16 widely used functional network atlases that are publicly available in fsaverage6 surface space, fs_LR_32k surface space, and MNI space including the Laird2011, Yeo2011, Shirer2012, Smith2013, Shen2013, Laumann2015, Miller2016, Gordon2016, Gordon2017, Schaefer2018, and Ji2019 ^14–25^. The majority of these atlases are derived from resting state fMRI data, though a few are based on task fMRI. Some of these atlases have multiple resolutions, resulting in 16 different atlases (**Supplementary Table 1**). We focused on cortical networks in this study; the non-cortical regions were masked for all atlases. Networks only corresponding to subcortical regions and the cerebellum were also excluded from all parcellations examined.

### Network similarity

To compute the network similarity between the input data and the atlases included in NCT, we treat the input data space as the reference data space and project the atlases to the reference atlas space (see section *Projection between atlases in different spaces* for projection details). In the case of input data being an atlas, the input data will be referred to as the “reference atlas”. **Figure 1** shows an example where the Yeo 17-network atlas in fsaverage6 space (the center) served as the reference atlas. All other surrounding atlases in different spaces were projected to the fsaverage6 space. The network similarity is defined as the Dice coefficients between the input data and different atlases in the reference data space.

### Spin tests

We use the spin test ^20,26,27^ implemented in Python ^28^ to assess if the reference input data significantly overlaps with networks from different atlases. Specifically, the reference input data is randomly rotated 1000 times. For each network, we compute the Dice coefficient between the rotated input data and this network. For a network highly overlapping with the input data, we expect the Dice coefficient between this network and the rotated input data to be smaller than the Dice coefficient between this network and the original input data. The *p*-value is defined as the number of rotated input data whose Dice coefficients are larger than the original input data divided by the total number of rotations. However, the spin test only works for data in surface space. For volumetric data, we project the data to fsaverage6 using an advanced nonlinear mapping approach ^39^ and perform the spin test in fsaverage6 surface space.

### Projection between atlases in different spaces

For atlases included in NCT, we treated each atlas as the reference atlas and projected other atlases to the reference atlas space.

1. Volume to volume projection. If the reference atlas and the other atlas were in different volumetric spaces, we used antsRegistration to register the other atlas template to the reference atlas template.
2. Volume to surface projection. If the reference atlas was in fsaverage6 surface space and the other atlas was in volumetric space, we first projected the other atlas into MNI152 volumetric space and then used an advanced nonlinear mapping approach ^39^ to project the other atlas from MNI152 to fsaverage6 surface space. If the reference atlas was in fs_LR_32k surface space ^40^ and the other atlas was in volumetric space, we first projected the other atlas into fsaverage6 surface space as above and then projected it from fsaverage6 surface space to fs_LR_32k surface space using the fsaverage-to-fs_LR registration provided by Human Connectome Project (HCP; https://wiki.humanconnectome.org/download/attachments/63078513/Resampling- FreeSurfer-HCP_5_8.pdf).
3. Surface to volume projection. If the reference atlas was in volumetric space and the other atlas was in fsaverage6 surface space, we first projected the other atlas to MNI152 space using an advanced nonlinear mapping approach ^39^ and then projected it from MNI152 to the reference volumetric template using antsRegistration. If the reference atlas was in volumetric space and the other atlas was in fs_LR_32k surface space, we first projected the other atlas to fsaverage6 surface space as above and then projected it from fsaverage6 to fs_LR_32k using the fsaverage-to-fs_LR registration provided by HCP.
4. Surface to surface projection. If the reference atlas was in fsaverage6 surface space and the other atlas was in fs_LR_32k surface space, we projected the other atlas to fsaverage6 surface space using the fs_LR-to-fsaverage registration provided by HCP. If the reference atlas was in fs_LR_32k surface space and the other atlas was in fsaverage6 surface space, we projected the other atlas to fs_LR_32k surface space using the fsaverage-to- fs_LR registration provided by HCP.

### HCP and UKBiobank data examples

HCP and UKB group ICA spatial maps were thresholded by fitting a spatial mixture model to the histogram of intensity values in each independent component spatial map. Mixture modeling explicitly models the null (background noise) and signal (“activated” or “deactivated”) parts of the spatial map. A threshold value of 0.6 was set to give slightly more importance to identifying signal over noise. In practice, the resulting thresholded spatial maps are robust to small deviations (e.g., threshold value = 0.5, which places equal importance on labeling a voxel as signal vs noise, and higher result in spatial maps that are very similar).

### Code availability

The *Network Correspondence Toolbox* is a Python toolbox that is available at the GitHub repository: https://github.com/rubykong/cbig_network_correspondence. The toolbox has been released to the Python Package Index (PyPI): https://pypi.org/project/cbig-network-correspondence. Detailed documentation and a tutorial are available at https://rubykong.github.io/cbig_network_correspondence.

## Acknowledgements

The authors would like to thank Taylor Bolt, Sarah M. Olshan, and Samar ElSayed for testing the *NCT* and providing feedback. They would also like to thank members of the OHBM Best Practices Committee for their support of this project. RK and BTTY are supported by the NUS Yong Loo Lin School of Medicine (NUHSRO/2020/124/TMR/LOA), the Singapore National Medical Research Council (NMRC) LCG (OFLCG19May-0035), NMRC CTG-IIT (CTGIIT23jan-0001), NMRC STaR (STaR20nov-0003), Singapore Ministry of Health (MOH) Centre Grant (CG21APR1009), the Temasek Foundation (TF2223-IMH-01), and the United States National Institutes of Health (R01MH120080 & R01MH133334). Any opinions, findings and conclusions or recommendations expressed in this material are those of the authors and do not reflect the views of the Singapore NMRC, MOH or Temasek Foundation. LQU is supported by R21HD111805 from NICHD and U01DA050987 from NIDA.

---

**Supplementary Table 1.** Sixteen widely used brain atlases in fsaverage6 surface space, fs_LR_32k surface space, and MNI space were included in the NCT. The first column shows the studies from which the atlases were obtained. The second column shows the specific atlas we included in the NCT.

| Abbreviation | Reference | Atlas |
| --- | --- | --- |
| Laird2011-20 | Laird2011 | Laird2011 20-node ICA-BrainMap maps |
| Yeo2011-7 | Yeo2011 | Yeo2011 7 networks |
| Yeo2011-17 | Yeo2011 | Yeo2011 17 networks |
| Shirer2012-90-14 | Shirer2012 | Shirer2012 90-ROI 14 networks |
| Smith2013-25 | Smith2013 | HCP 25-node ICA maps |
| Shen2013-268-8 | Shen2013 | Shen2013 268-ROI with 8 networks |
| Shen2013-368-8 | Shen2013 | Shen2013 368-ROI with 8 networks |
| Laumann2015-12 | Laumann2015 | Laumann2015 12 networks (Power2011) |
| Miller2016-25 | Miller2016 | UK Biobank 25-node ICA maps |
| Gordon2016-286-12 | Gordon2016 | Gordon2016 286-ROI with 12 networks |
| Gordon2017-17 | Gordon2017 | Gordon2017 17 networks |
| Schaefer2018-200+Kong2021-17 | Schaefer2018 | Schaefer2018 200-ROI with Kong2021 17 networks |
| Schaefer2018-200+Yeo2011-17 | Schaefer2018 | Schaefer2018 200-ROI with Yeo2011 17 networks |
| Schaefer2018-400+Kong2021-17 | Schaefer2018 | Schaefer2018 400-ROI with Kong2021 17 networks |
| Schaefer2018-400+Yeo2011-17 | Schaefer2018 | Schaefer2018 400-ROI with Yeo2011 17 networks |
| Glasser2016-360+Ji2019-12 | Ji2019 | Glasser2016 360-ROI with Ji2019 12 Cole-Anticevic networks |

**Supplementary Table 2.** Network-to-network spatial correspondence between all 136 network pairs from the Yeo2011-17 (rows) and Gordon2017-17 (columns) atlases (Dice coefficients and corresponding *p*-values).

|  | **Default** | **LatVis** | **FrontPar** | **MedVis** | **DorsAttn** | **Premotor** | **Language** | **Salience** | **CingOperc** |
| --- | --- | --- | --- | --- | --- | --- | --- | --- | --- |
| **VisualA** | 0.0000<br>(p=0.9461) | 0.7311<br>(p=0.0010) | 0.0000<br>(p=0.9131) | 0.0858<br>(p=0.1548) | 0.0682<br>(p=0.3437) | 0.0000<br>(p=0.5225) | 0.0000<br>(p=0.6763) | 0.0000<br>(p=0.4665) | 0.0000<br>(p=0.9251) |
| **VisualB** | 0.0005<br>(p=0.9191) | 0.3164<br>(p=0.0799) | 0.0000<br>(p=0.9091) | 0.6462<br>(p=0.0030) | 0.0004<br>(p=0.6993) | 0.0000<br>(p=0.5145) | 0.0000<br>(p=0.6593) | 0.0000<br>(p=0.4515) | 0.0000<br>(p=0.9081) |
| **SomatomotorA** | 0.0000<br>(p=0.9860) | 0.0000<br>(p=0.6523) | 0.0000<br>(p=0.9790) | 0.0000<br>(p=0.3566) | 0.0000<br>(p=0.8821) | 0.0738<br>(p=0.2318) | 0.0000<br>(p=0.8272) | 0.0000<br>(p=0.5694) | 0.0207<br>(p=0.7512) |
| **SomatomotorB** | 0.0000<br>(p=0.9860) | 0.0000<br>(p=0.5814) | 0.0000<br>(p=0.9680) | 0.0000<br>(p=0.3536) | 0.0000<br>(p=0.8122) | 0.0108<br>(p=0.4076) | 0.0098<br>(p=0.6384) | 0.0000<br>(p=0.5415) | 0.0759<br>(p=0.4605) |
| **DorsAttnA** | 0.0011<br>(p=0.9860) | 0.1205<br>(p=0.2198) | 0.0307<br>(p=0.6444) | 0.0000<br>(p=0.4336) | 0.6204<br>(p=0.0010) | 0.0098<br>(p=0.4965) | 0.0222<br>(p=0.5534) | 0.0000<br>(p=0.6434) | 0.0011<br>(p=0.9580) |
| **DorsAttnB** | 0.0000<br>(p=1.0000) | 0.0000<br>(p=0.6973) | 0.0059<br>(p=0.9321) | 0.0000<br>(p=0.4595) | 0.2730<br>(p=0.0010) | 0.6132<br>(p=0.0010) | 0.0101<br>(p=0.7303) | 0.0000<br>(p=0.6364) | 0.0736<br>(p=0.4715) |
| **Sal/VenAttnA** | 0.0004<br>(p=1.0000) | 0.0000<br>(p=0.8731) | 0.0031<br>(p=0.9750) | 0.0000<br>(p=0.6054) | 0.0018<br>(p=0.9590) | 0.0114<br>(p=0.6543) | 0.0099<br>(p=0.8312) | 0.0120<br>(p=0.6134) | 0.7310<br>(p=0.0010) |
| **Sal/VenAttnB** | 0.0022<br>(p=0.9970) | 0.0000<br>(p=0.8172) | 0.1058<br>(p=0.2078) | 0.0000<br>(p=0.5724) | 0.0063<br>(p=0.8631) | 0.0000<br>(p=0.7912) | 0.0556<br>(p=0.3676) | 0.3719<br>(p=0.0020) | 0.2913<br>(p=0.0010) |
| **LimbicA** | 0.0162<br>(p=0.6653) | 0.0029<br>(p=0.3237) | 0.0354<br>(p=0.4066) | 0.0000<br>(p=0.2038) | 0.0009<br>(p=0.5734) | 0.0000<br>(p=0.4266) | 0.0060<br>(p=0.4915) | 0.0000<br>(p=0.3636) | 0.0127<br>(p=0.5634) |
| **LimbicB** | 0.0879<br>(p=0.2078) | 0.0002<br>(p=0.3676) | 0.0757<br>(p=0.2008) | 0.0000<br>(p=0.2068) | 0.0000<br>(p=0.5914) | 0.0000<br>(p=0.4106) | 0.0000<br>(p=0.5075) | 0.0004<br>(p=0.3247) | 0.0095<br>(p=0.5205) |
| **ControlC** | 0.0796<br>(p=0.1658) | 0.0000<br>(p=0.4815) | 0.0036<br>(p=0.7133) | 0.0000<br>(p=0.2517) | 0.0229<br>(p=0.4476) | 0.0000<br>(p=0.5135) | 0.0000<br>(p=0.6793) | 0.0000<br>(p=0.4256) | 0.0296<br>(p=0.4835) |
| **ControlA** | 0.0026<br>(p=0.9960) | 0.0000<br>(p=0.9271) | 0.5897<br>(p=0.0010) | 0.0000<br>(p=0.6424) | 0.1430<br>(p=0.0889) | 0.0000<br>(p=0.8901) | 0.0179<br>(p=0.7622) | 0.0000<br>(p=0.8222) | 0.0066<br>(p=0.9770) |
| **ControlB** | 0.2171<br>(p=0.0050) | 0.0000<br>(p=0.8981) | 0.3880<br>(p=0.0010) | 0.0000<br>(p=0.5824) | 0.0052<br>(p=0.9041) | 0.0000<br>(p=0.8751) | 0.0202<br>(p=0.6993) | 0.0497<br>(p=0.2697) | 0.0029<br>(p=0.9900) |
| **TempPar** | 0.0061<br>(p=0.8521) | 0.0000<br>(p=0.6474) | 0.0000<br>(p=0.9381) | 0.0000<br>(p=0.3716) | 0.0000<br>(p=0.8182) | 0.0000<br>(p=0.5754) | 0.6375<br>(p=0.0010) | 0.0000<br>(p=0.5385) | 0.0091<br>(p=0.6513) |
| **DefaultC** | 0.0518<br>(p=0.3267) | 0.0024<br>(p=0.4765) | 0.0013<br>(p=0.8651) | 0.0255<br>(p=0.2178) | 0.0003<br>(p=0.7902) | 0.0000<br>(p=0.5774) | 0.0090<br>(p=0.5235) | 0.0000<br>(p=0.5055) | 0.0004<br>(p=0.9381) |
| **DefaultA** | 0.5622<br>(p=0.0010) | 0.0000<br>(p=0.9161) | 0.0044<br>(p=0.9660) | 0.0000<br>(p=0.5984) | 0.0000<br>(p=0.9830) | 0.0000<br>(p=0.9181) | 0.0149<br>(p=0.7682) | 0.0786<br>(p=0.1648) | 0.0000<br>(p=0.9990) |
| **DefaultB** | 0.3694<br>(p=0.0010) | 0.0000<br>(p=0.8641) | 0.0193<br>(p=0.8901) | 0.0000<br>(p=0.6004) | 0.0003<br>(p=0.9680) | 0.0000<br>(p=0.8392) | 0.3259<br>(p=0.0020) | 0.0831<br>(p=0.1449) | 0.0032<br>(p=0.9830) |

**Supplementary Table 2 (cont.).** Network-to-network spatial correspondence between all 136 network pairs from the Yeo2011-17 (rows) and Gordon2017-17 (columns) atlases (Dice coefficients and corresponding *p*-values).
|  | **HandSM** | **FaceSM** | **Auditory** | **AntMTL** | **PostMTL** | **ParMemory** | **Context** | **FootSM** |
| --- | --- | --- | --- | --- | --- | --- | --- | --- |
| **VisualA** | 0.0000<br>(p=0.5994) | 0.0000<br>(p=0.3287) | 0.0000<br>(p=0.4116) | 0.0000<br>(p=0.2637) | 0.0000<br>(p=0.2408) | 0.0000<br>(p=0.2777) | 0.0043<br>(p=0.4565) | 0.0000<br>(p=0.3217) |
| **VisualB** | 0.0000<br>(p=0.5944) | 0.0000<br>(p=0.3137) | 0.0000<br>(p=0.4196) | 0.0000<br>(p=0.2517) | 0.0030<br>(p=0.1718) | 0.0310<br>(p=0.1898) | 0.0635<br>(p=0.2058) | 0.0000<br>(p=0.3277) |
| **SomatomotorA** | 0.5121<br>(p=0.0010) | 0.0332<br>(p=0.2258) | 0.0161<br>(p=0.3197) | 0.0000<br>(p=0.4056) | 0.0000<br>(p=0.3127) | 0.0000<br>(p=0.3636) | 0.0000<br>(p=0.7043) | 0.6892<br>(p=0.0010) |
| **SomatomotorB** | 0.0076<br>(p=0.4875) | 0.4738<br>(p=0.0020) | 0.6948<br>(p=0.0010) | 0.0000<br>(p=0.3487) | 0.0000<br>(p=0.2987) | 0.0000<br>(p=0.3516) | 0.0000<br>(p=0.6883) | 0.0000<br>(p=0.3716) |
| **DorsAttnA** | 0.0016<br>(p=0.7223) | 0.0000<br>(p=0.4885) | 0.0000<br>(p=0.6454) | 0.0000<br>(p=0.4665) | 0.0000<br>(p=0.3596) | 0.0186<br>(p=0.2787) | 0.0496<br>(p=0.2617) | 0.0000<br>(p=0.5195) |
| **DorsAttnB** | 0.0543<br>(p=0.2697) | 0.0102<br>(p=0.3956) | 0.0000<br>(p=0.6783) | 0.0000<br>(p=0.4815) | 0.0000<br>(p=0.3916) | 0.0000<br>(p=0.4246) | 0.0000<br>(p=0.7652) | 0.0012<br>(p=0.5175) |
| **Sal/VenAttnA** | 0.0114<br>(p=0.6194) | 0.0094<br>(p=0.4705) | 0.0213<br>(p=0.4855) | 0.0023<br>(p=0.5734) | 0.0000<br>(p=0.5175) | 0.0027<br>(p=0.5185) | 0.0000<br>(p=0.8801) | 0.0100<br>(p=0.5085) |
| **Sal/VenAttnB** | 0.0000<br>(p=0.8781) | 0.0000<br>(p=0.5774) | 0.0000<br>(p=0.7223) | 0.0000<br>(p=0.6014) | 0.0000<br>(p=0.4735) | 0.0063<br>(p=0.4346) | 0.0000<br>(p=0.8482) | 0.0000<br>(p=0.6394) |
| **LimbicA** | 0.0237<br>(p=0.2498) | 0.0000<br>(p=0.2148) | 0.0000<br>(p=0.3457) | 0.7469<br>(p=0.0010) | 0.0183<br>(p=0.1159) | 0.0000<br>(p=0.2148) | 0.0000<br>(p=0.4695) | 0.0000<br>(p=0.2488) |
| **LimbicB** | 0.0000<br>(p=0.4446) | 0.0000<br>(p=0.2338) | 0.0000<br>(p=0.3087) | 0.0000<br>(p=0.1988) | 0.0000<br>(p=0.1538) | 0.0000<br>(p=0.1838) | 0.0000<br>(p=0.4086) | 0.0000<br>(p=0.2398) |
| **ControlC** | 0.0000<br>(p=0.5325) | 0.0000<br>(p=0.3237) | 0.0000<br>(p=0.4096) | 0.0000<br>(p=0.2727) | 0.0000<br>(p=0.1858) | 0.4955<br>(p=0.0010) | 0.0305<br>(p=0.2597) | 0.0000<br>(p=0.3397) |
| **ControlA** | 0.0000<br>(p=0.9201) | 0.0003<br>(p=0.6484) | 0.0000<br>(p=0.8072) | 0.0000<br>(p=0.6703) | 0.0000<br>(p=0.5375) | 0.0000<br>(p=0.6114) | 0.0031<br>(p=0.7632) | 0.0000<br>(p=0.7502) |
| **ControlB** | 0.0000<br>(p=0.8851) | 0.0000<br>(p=0.6224) | 0.0000<br>(p=0.7652) | 0.0000<br>(p=0.6553) | 0.0000<br>(p=0.5205) | 0.0000<br>(p=0.5784) | 0.0069<br>(p=0.6903) | 0.0000<br>(p=0.7123) |
| **TempPar** | 0.0025<br>(p=0.5305) | 0.0000<br>(p=0.3576) | 0.0358<br>(p=0.1748) | 0.0000<br>(p=0.4176) | 0.0000<br>(p=0.2747) | 0.0000<br>(p=0.3307) | 0.0000<br>(p=0.5844) | 0.0000<br>(p=0.4595) |
| **DefaultC** | 0.0000<br>(p=0.6384) | 0.0000<br>(p=0.3776) | 0.0000<br>(p=0.4725) | 0.0253<br>(p=0.2108) | 0.0661<br>(p=0.0340) | 0.0000<br>(p=0.3327) | 0.6406<br>(p=0.0010) | 0.0000<br>(p=0.3996) |
| **DefaultA** | 0.0000<br>(p=0.8981) | 0.0000<br>(p=0.6783) | 0.0000<br>(p=0.8062) | 0.0000<br>(p=0.6643) | 0.0000<br>(p=0.4975) | 0.0543<br>(p=0.2068) | 0.0425<br>(p=0.3227) | 0.0000<br>(p=0.7612) |
| **DefaultB** | 0.0000<br>(p=0.8911) | 0.0000<br>(p=0.6294) | 0.0000<br>(p=0.7672) | 0.0309<br>(p=0.3896) | 0.0000<br>(p=0.5235) | 0.0038<br>(p=0.5095) | 0.0000<br>(p=0.8721) | 0.0000<br>(p=0.6843) |

**Supplementary Table 3.** Network spatial correspondence between HCP working memory task contrast (2BK vs. 0BK) and networks from 8 atlases (Dice coefficient and corresponding *p*- values).

| Glasser2016-360+Ji2019-12 |  |  | Schaefer2018-400+Kong2019-17 |  |  | Shirer2013-90-14 |  |  |
| --- | --- | --- | --- | --- | --- | --- | --- | --- |
| Name | Dice | P value | Name | Dice | P value | Name | Dice | P value |
| Visual1 | 0.0031 | 0.7423 | DefaultA | 0.023 | 0.9161 | AntSal | 0.1876 | 0.009* |
| Visual2 | 0.0072 | 0.8911 | DefaultB | 0.0394 | 0.8102 | Auditory | 0.0001 | 0.7762 |
| Somatomotor | 0.0001 | 0.997 | DefaultC | 0.1006 | 0.2308 | DorsalDMN | 0.002 | 0.9091 |
| CingOperc | 0.225 | 0.0999 | Language | 0.0583 | 0.4795 | HighVisual | 0.0001 | 0.7183 |
| DorsAttn | 0.2619 | 0.005* | ControlA | 0.3119 | 0.001* | Language | 0.0049 | 0.7473 |
| Language | 0.0455 | 0.6304 | ControlB | 0.2559 | 0.001* | LECN | 0.0874 | 0.042* |
| FrontPar | 0.4751 | 0.001* | ControlC | 0.1461 | 0.032* | PostSal | 0.0521 | 0.2348 |
| Auditory | 0.0001 | 0.7253 | Sal/VenAttnA | 0.0388 | 0.7852 | Precuneus | 0.078 | 0.1149 |
| Default | 0.0561 | 0.9381 | Sal/VenAttnB | 0.1546 | 0.0979 | PrimVisual | 0.0001 | 0.5664 |
| PostMulti | 0.0251 | 0.4016 | DorsAttnA | 0.221 | 0.011* | RECN | 0.2364 | 0.001* |
| VentMulti | 0.0001 | 0.6593 | DorsAttnB | 0.0406 | 0.6024 | Sensorimotor | 0.0001 | 0.8352 |
| OrbitAffective | 0.0006 | 0.5904 | Auditory | 0.0001 | 0.9111 | VentralDMN | 0.1819 | 0.021* |
| - | - | - | SomatomotorA | 0.0001 | 0.952 | Visuospatial | 0.1994 | 0.041* |
| - | - | - | SomatomotorB | 0.0003 | 0.9191 | - | - | - |
| - | - | - | VisualA | 0.0289 | 0.7782 | - | - | - |
| - | - | - | VisualB | 0.0032 | 0.8641 | - | - | - |
| - | - | - | VisualC | 0.0018 | 0.5604 | - | - | - |

| Yeo2011-17 |  |  | Shen2013-268-8 |  |  | Laird2011-20 |  |  |
| --- | --- | --- | --- | --- | --- | --- | --- | --- |
| Name | Dice | P value | Name | Dice | P value | Name | Dice | P value |
| VisualA | 0.0027 | 0.8262 | MedFront | 0.1151 | 0.5365 | Emo/Interoception1 | 0.0139 | 0.9331 |
| VisualB | 0.002 | 0.7942 | FrontPar | 0.3676 | 0.007* | Emo/Interoception2 | 0.104 | 0.6004 |
| SomatomotorA | 0.0001 | 0.958 | Default | 0.0495 | 0.7383 | Emo/Interoception3 | 0.0524 | 0.5594 |
| SomatomotorB | 0.0001 | 0.9341 | Motor | 0.0701 | 0.8981 | Emo/Interoception4 | 0.1775 | 0.3876 |
| DorsAttnA | 0.097 | 0.3277 | VisualA | 0.028 | 0.6783 | Mot/Visspatial1 | 0.2556 | 0.1489 |
| DorsAttnB | 0.1206 | 0.2507 | VisualB | 0.0055 | 0.5844 | Mot/Visspatial2 | 0.4077 | 0.003* |
| Sal/VenAttnA | 0.0261 | 0.9051 | VisAssoc | 0.1388 | 0.3337 | Mot/Visspatial3 | 0.1743 | 0.4575 |
| Sal/VenAttnB | 0.2699 | 0.001* | SalSubcor | 0.2523 | 0.025* | Mot/Visspatial4 | 0.0404 | 0.9131 |
| LimbicA | 0.0001 | 0.7972 | - | - | - | Visual1 | 0.0794 | 0.8881 |
| LimbicB | 0.0023 | 0.6633 | - | - | - | Visual2 | 0.0398 | 0.6793 |
| ControlC | 0.0986 | 0.1249 | - | - | - | Visual3 | 0.0508 | 0.7103 |
| ControlA | 0.34 | 0.001* | - | - | - | DivergentCog1 | 0.1651 | 0.6184 |
| ControlB | 0.3622 | 0.001* | - | - | - | DivergentCog3 | 0.3949 | 0.005* |
| TempPar | 0.0067 | 0.7103 | - | - | - | DivergentCog4 | 0.0423 | 0.8631 |
| DefaultC | 0.0077 | 0.7003 | - | - | - | DivergentCog5 | 0.1287 | 0.5544 |
| DefaultA | 0.0534 | 0.7093 | - | - | - | DivergentCog6 | 0.3031 | 0.001* |
| DefaultB | 0.037 | 0.8102 | - | - | - | - | - | - |

**Supplementary Table 3 (Cont.).** Network spatial correspondence between HCP working memory task contrast (2BK vs. 0BK) and networks from 8 atlases (Dice coefficient and corresponding *p*-values).
| Laumann2015-12 |  |  | Gordon2017-17 |  |  |
| --- | --- | --- | --- | --- | --- |
| Name | Dice | P value | Name | Dice | P value |
| Default | 0.094 | 0.6364 | Default | 0.0972 | 0.6953 |
| Visual | 0.2123 | 0.2228 | LatVis | 0.2355 | 0.1638 |
| FrontPar | 0.1293 | 0.2677 | FrontPar | 0.1321 | 0.3087 |
| DorsAttn | 0.1371 | 0.2697 | MedVis | 0.0024 | 0.6743 |
| VentAttn | 0.3729 | 0.002* | DorsAttn | 0.1873 | 0.1299 |
| Salience | 0.0001 | 0.7253 | Premotor | 0.0004 | 0.9201 |
| CingOperc | 0.0344 | 0.8671 | Language | 0.3926 | 0.002* |
| SM | 0.0001 | 0.981 | Salience | 0.0039 | 0.7732 |
| LateralSM | 0.0001 | 0.7922 | CingOperc | 0.0329 | 0.8781 |
| Auditory | 0.0004 | 0.8332 | HandSM | 0.0001 | 0.97 |
| MedPar | 0.0001 | 0.7223 | FaceSM | 0.0001 | 0.7552 |
| ParOcc | 0.1264 | 0.014* | Auditory | 0.0007 | 0.8591 |
| - | - | - | AntMTL | 0.0329 | 0.3536 |
| - | - | - | PostMTL | 0.0007 | 0.5594 |
| - | - | - | ParMemory | 0.0001 | 0.7283 |
| - | - | - | Context | 0.1473 | 0.029* |
| - | - | - | FootSM | 0.0001 | 0.8651 |

**Supplementary Table 4.**
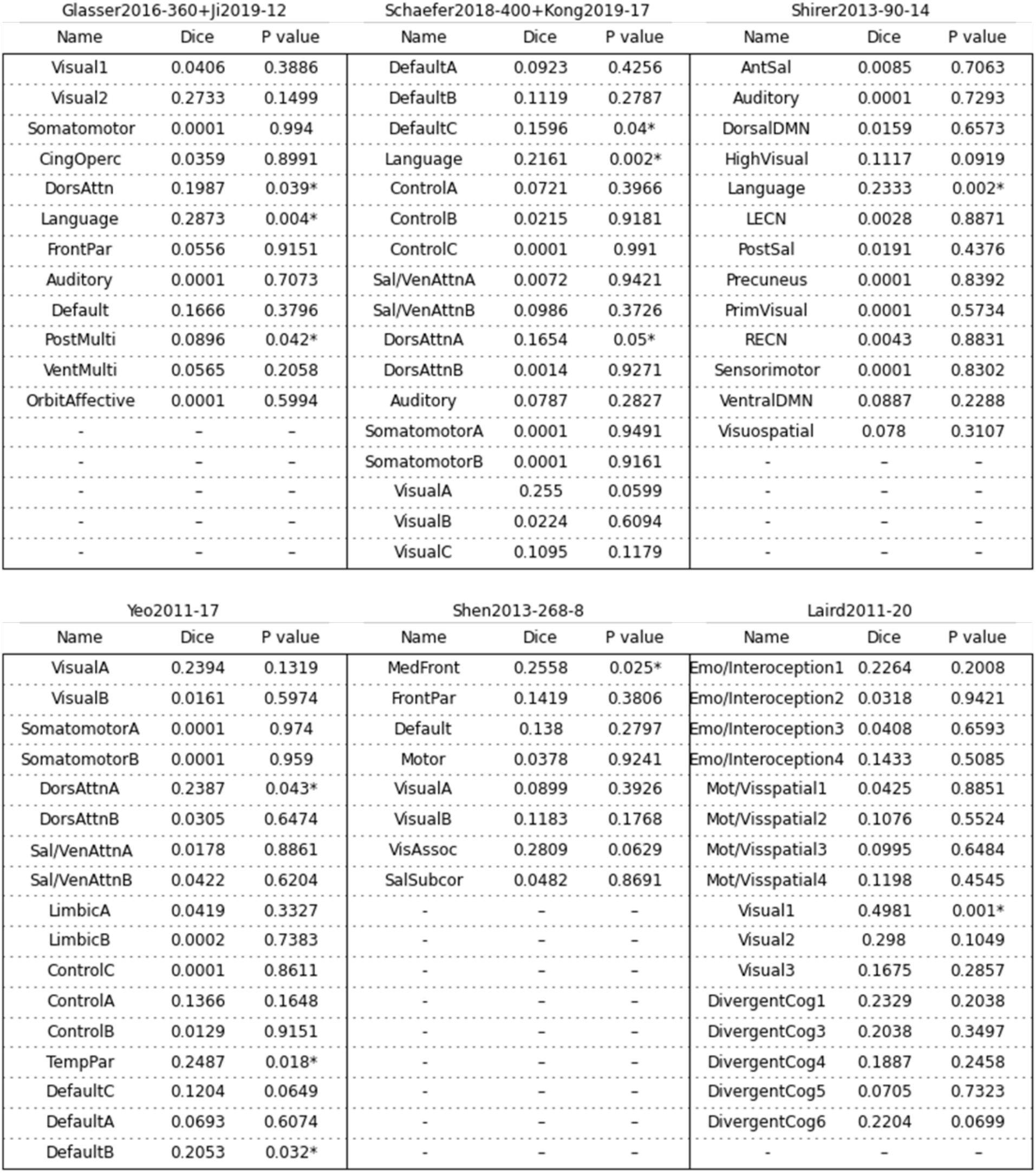
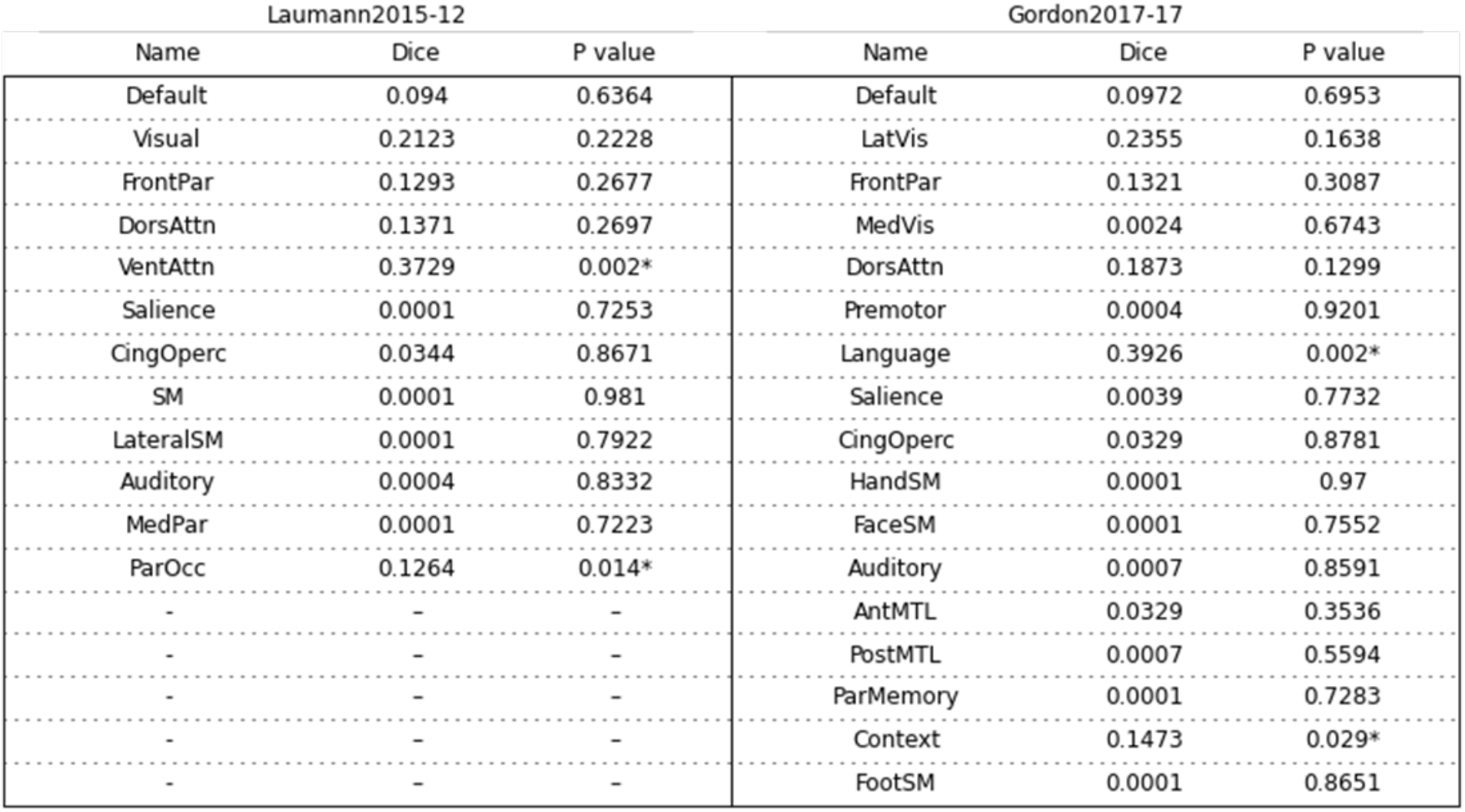
Network spatial correspondence between HCP social task contrast (theory of mind vs. random) and networks from 8 atlases (Dice coefficient and corresponding *p*- values).

**Supplementary Table 5.** Network spatial correspondence between UKB ICA component 5 and networks from 8 atlases (Dice coefficient and corresponding *p*-values).

| Glasser2016-360+Ji2019-12 |  |  | Schaefer2018-400+Kong2019-17 |  |  | Shirer2013-90-14 |  |  |
| --- | --- | --- | --- | --- | --- | --- | --- | --- |
| Name | Dice | P value | Name | Dice | P value | Name | Dice | P value |
| Visual1 | 0.2671 | 0.003* | DefaultA | 0.0135 | 0.984 | AntSal | 0.0556 | 0.5415 |
| Visual2 | 0.3304 | 0.0549 | DefaultB | 0.0364 | 0.969 | Auditory | 0.012 | 0.6094 |
| Somatomotor | 0.2149 | 0.3966 | DefaultC | 0.1202 | 0.1878 | DorsalDMN | 0.0255 | 0.8412 |
| CingOperc | 0.1867 | 0.4745 | Language | 0.0128 | 0.97 | HighVisual | 0.0057 | 0.5994 |
| DorsAttn | 0.0375 | 0.8831 | ControlA | 0.0374 | 0.9471 | Language | 0.0032 | 0.9131 |
| Language | 0.0285 | 0.8951 | ControlB | 0.0162 | 0.998 | LECN | 0.0054 | 0.9421 |
| FrontPar | 0.0858 | 0.952 | ControlC | 0.0594 | 0.6603 | PostSal | 0.0116 | 0.7063 |
| Auditory | 0.0538 | 0.3357 | Sal/VenAttnA | 0.1218 | 0.4186 | Precuneus | 0.0325 | 0.4795 |
| Default | 0.087 | 0.9101 | Sal/VenAttnB | 0.0727 | 0.6943 | PrimVisual | 0.0701 | 0.004* |
| PostMulti | 0.0259 | 0.4535 | DorsAttnA | 0.0341 | 0.9091 | RECN | 0.0076 | 0.8951 |
| VentMulti | 0.0001 | 0.6873 | DorsAttnB | 0.0397 | 0.7712 | Sensorimotor | 0.087 | 0.2138 |
| OrbitAffective | 0.0057 | 0.6494 | Auditory | 0.0598 | 0.4755 | VentralDMN | 0.0938 | 0.2058 |
| - | - | - | SomatomotorA | 0.092 | 0.4835 | Visuospatial | 0.0075 | 0.981 |
| - | - | - | SomatomotorB | 0.1627 | 0.1808 | - | - | - |
| - | - | - | VisualA | 0.1709 | 0.1788 | - | - | - |
| - | - | - | VisualB | 0.4559 | 0.001* | - | - | - |
| - | - | - | VisualC | 0.0447 | 0.2058 | - | - | - |

| Yeo2011-17 |  |  | Shen2013-268-8 |  |  | Laird2011-20 |  |  |
| --- | --- | --- | --- | --- | --- | --- | --- | --- |
| Name | Dice | P value | Name | Dice | P value | Name | Dice | P value |
| VisualA | 0.2016 | 0.1029 | MedFront | 0.0727 | 0.9331 | Emo/Interoception1 | 0.0705 | 0.9201 |
| VisualB | 0.4076 | 0.001* | FrontPar | 0.0155 | 0.996 | Emo/Interoception2 | 0.0923 | 0.8801 |
| SomatomotorA | 0.1494 | 0.3776 | Default | 0.0897 | 0.5395 | Emo/Interoception3 | 0.0552 | 0.7213 |
| SomatomotorB | 0.1535 | 0.3027 | Motor | 0.2258 | 0.5564 | Emo/Interoception4 | 0.1537 | 0.8791 |
| DorsAttnA | 0.0671 | 0.6234 | VisualA | 0.4734 | 0.001* | Mot/Visspatial1 | 0.2776 | 0.2388 |
| DorsAttnB | 0.0642 | 0.6154 | VisualB | 0.1176 | 0.1109 | Mot/Visspatial2 | 0.2667 | 0.2767 |
| Sal/VenAttnA | 0.1453 | 0.2927 | VisAssoc | 0.0645 | 0.7423 | Mot/Visspatial3 | 0.2032 | 0.5774 |
| Sal/VenAttnB | 0.0882 | 0.5834 | SalSubcor | 0.2016 | 0.4476 | Mot/Visspatial4 | 0.1933 | 0.3317 |
| LimbicA | 0.0001 | 0.8142 | - | - | - | Visual1 | 0.2736 | 0.2178 |
| LimbicB | 0.0001 | 0.7832 | - | - | - | Visual2 | 0.2209 | 0.1139 |
| ControlC | 0.0528 | 0.4176 | - | - | - | Visual3 | 0.5793 | 0.001* |
| ControlA | 0.0185 | 0.985 | - | - | - | DivergentCog1 | 0.2408 | 0.4655 |
| ControlB | 0.019 | 0.972 | - | - | - | DivergentCog3 | 0.0883 | 0.957 |
| TempPar | 0.003 | 0.8861 | - | - | - | DivergentCog4 | 0.1039 | 0.8242 |
| DefaultC | 0.0913 | 0.1069 | - | - | - | DivergentCog5 | 0.1916 | 0.4575 |
| DefaultA | 0.0274 | 0.9271 | - | - | - | DivergentCog6 | 0.1137 | 0.9301 |
| DefaultB | 0.0456 | 0.9451 | - | - | - | - | - | - |

**Supplementary Table 5 (Cont.).** Network spatial correspondence between UKB ICA component 5 and networks from 8 atlases (Dice coefficient and corresponding *p*-values).
| Laumann2015-12 |  |  | Gordon2017-17 |  |  |
| --- | --- | --- | --- | --- | --- |
| Name | Dice | P value | Name | Dice | P value |
| Default | 0.0602 | 0.9371 | Default | 0.0452 | 0.998 |
| Visual | 0.5032 | 0.005* | LatVis | 0.3346 | 0.041* |
| FrontPar | 0.0345 | 0.9401 | FrontPar | 0.0261 | 0.995 |
| DorsAttn | 0.0553 | 0.8801 | MedVis | 0.2568 | 0.002* |
| VentAttn | 0.0144 | 0.962 | DorsAttn | 0.056 | 0.8192 |
| Saliency | 0.0161 | 0.4995 | Premotor | 0.0245 | 0.7742 |
| CingOperc | 0.2082 | 0.2967 | Language | 0.0244 | 0.9221 |
| SM | 0.1602 | 0.3826 | Saliency | 0.0624 | 0.4615 |
| LateralSM | 0.08 | 0.2118 | CingOperc | 0.1908 | 0.3477 |
| Auditory | 0.0835 | 0.3646 | HandSM | 0.0873 | 0.2927 |
| MedPar | 0.0322 | 0.2997 | FaceSM | 0.0775 | 0.1788 |
| ParOcc | 0.0725 | 0.2238 | Auditory | 0.1129 | 0.2957 |
| - | - | - | AntMTL | 0.0001 | 0.8312 |
| - | - | - | PostMTL | 0.0059 | 0.6833 |
| - | - | - | ParMemory | 0.0364 | 0.3337 |
| - | - | - | Context | 0.1215 | 0.0519 |
| - | - | - | FootSM | 0.0741 | 0.4086 |

**Supplementary Table 6.** Network spatial correspondence between UKB ICA component 3 and networks from 8 atlases (Dice coefficient and corresponding *p*-values).

| Glasser2016-360+Ji2019-12 |  |  | Schaefer2018-400+Kong2019-17 |  |  | Shirer2013-90-14 |  |  |
| --- | --- | --- | --- | --- | --- | --- | --- | --- |
| Name | Dice | P value | Name | Dice | P value | Name | Dice | P value |
| Visual1 | 0.0168 | 0.9061 | DefaultA | 0.0492 | 0.991 | AntSal | 0.061 | 0.5115 |
| Visual2 | 0.107 | 0.7433 | DefaultB | 0.0938 | 0.8472 | Auditory | 0.0387 | 0.3337 |
| Somatomotor | 0.2585 | 0.3696 | DefaultC | 0.0532 | 0.8811 | DorsalDMN | 0.0756 | 0.5315 |
| CingOperc | 0.4221 | 0.001* | Language | 0.0303 | 0.999 | HighVisual | 0.0016 | 0.9321 |
| DorsAttn | 0.2233 | 0.001* | ControlA | 0.1376 | 0.045* | Language | 0.0284 | 0.7552 |
| Language | 0.0383 | 0.993 | ControlB | 0.0612 | 0.992 | LECN | 0.0243 | 0.9121 |
| FrontPar | 0.203 | 0.7093 | ControlC | 0.0752 | 0.8092 | PostSal | 0.0844 | 0.001* |
| Auditory | 0.0324 | 0.8581 | Sal/VenAttnA | 0.2207 | 0.001* | Precuneus | 0.0388 | 0.6214 |
| Default | 0.1409 | 0.99 | Sal/VenAttnB | 0.2327 | 0.001* | PrimVisual | 0.0016 | 0.8751 |
| PostMulti | 0.0456 | 0.1199 | DorsAttnA | 0.2225 | 0.001* | RECN | 0.0367 | 0.6983 |
| VentMulti | 0.0381 | 0.6014 | DorsAttnB | 0.1886 | 0.001* | Sensorimotor | 0.0114 | 0.9371 |
| OrbitAffective | 0.0291 | 0.4615 | Auditory | 0.0646 | 0.7692 | VentralDMN | 0.0948 | 0.2028 |
| - | - | - | SomatomotorA | 0.0818 | 0.7702 | Visuospatial | 0.1842 | 0.001* |
| - | - | - | SomatomotorB | 0.0718 | 0.6863 | - | - | - |
| - | - | - | VisualA | 0.126 | 0.3307 | - | - | - |
| - | - | - | VisualB | 0.0223 | 0.9351 | - | - | - |
| - | - | - | VisualC | 0.0019 | 0.9121 | - | - | - |

| Yeo2011-17 |  |  | Shen2013-268-8 |  |  | Laird2011-20 |  |  |
| --- | --- | --- | --- | --- | --- | --- | --- | --- |
| Name | Dice | P value | Name | Dice | P value | Name | Dice | P value |
| VisualA | 0.0145 | 0.973 | MedFront | 0.1502 | 0.9371 | Emo/Interoception1 | 0.2306 | 0.6573 |
| VisualB | 0.011 | 0.97 | FrontPar | 0.2351 | 0.3417 | Emo/Interoception2 | 0.2422 | 0.5065 |
| SomatomotorA | 0.1199 | 0.7203 | Default | 0.1019 | 0.8012 | Emo/Interoception3 | 0.121 | 0.2807 |
| SomatomotorB | 0.1278 | 0.6693 | Motor | 0.4274 | 0.0779 | Emo/Interoception4 | 0.3903 | 0.0579 |
| DorsAttnA | 0.2025 | 0.002* | VisualA | 0.0302 | 0.963 | Mot/Visspatial1 | 0.3045 | 0.1908 |
| DorsAttnB | 0.2177 | 0.001* | VisualB | 0.0044 | 0.962 | Mot/Visspatial2 | 0.428 | 0.005* |
| Sal/VenAttnA | 0.2843 | 0.001* | VisAssoc | 0.2464 | 0.022* | Mot/Visspatial3 | 0.4858 | 0.001* |
| Sal/VenAttnB | 0.1447 | 0.1369 | SalSubcor | 0.2676 | 0.0539 | Mot/Visspatial4 | 0.2043 | 0.5654 |
| LimbicA | 0.0504 | 0.7602 | - | - | - | Visual1 | 0.3653 | 0.1808 |
| LimbicB | 0.0383 | 0.7083 | - | - | - | Visual2 | 0.085 | 0.8022 |
| ControlC | 0.0683 | 0.3956 | - | - | - | Visual3 | 0.0767 | 0.9411 |
| ControlA | 0.1944 | 0.005* | - | - | - | DivergentCog1 | 0.3851 | 0.5185 |
| ControlB | 0.0581 | 0.984 | - | - | - | DivergentCog3 | 0.3997 | 0.0929 |
| TempPar | 0.0277 | 0.7692 | - | - | - | DivergentCog4 | 0.2575 | 0.4805 |
| DefaultC | 0.024 | 0.7283 | - | - | - | DivergentCog5 | 0.3691 | 0.0589 |
| DefaultA | 0.0732 | 0.971 | - | - | - | DivergentCog6 | 0.2907 | 0.0649 |
| DefaultB | 0.0853 | 0.9201 | - | - | - | - | - | - |

**Supplementary Table 6 (Cont.).** Network spatial correspondence between UKB ICA component 3 and networks from 8 atlases (Dice coefficient and corresponding *p*-values).
| Laumann2015-12 |  |  | Gordon2017-17 |  |  |
| --- | --- | --- | --- | --- | --- |
| Name | Dice | P value | Name | Dice | P value |
| Default | 0.1147 | 0.986 | Default | 0.1273 | 0.988 |
| Visual | 0.0438 | 0.971 | LatVis | 0.035 | 0.9391 |
| FrontPar | 0.1773 | 0.1618 | FrontPar | 0.2354 | 0.0759 |
| DorsAttn | 0.312 | 0.001* | MedVis | 0.0105 | 0.9401 |
| VentAttn | 0.0491 | 0.8811 | DorsAttn | 0.263 | 0.001* |
| Saliency | 0.0132 | 0.5974 | Premotor | 0.141 | 0.001* |
| CingOperc | 0.3675 | 0.001* | Language | 0.0586 | 0.8801 |
| SM | 0.1569 | 0.5574 | Saliency | 0.0428 | 0.7952 |
| LateralSM | 0.0539 | 0.5734 | CingOperc | 0.3867 | 0.001* |
| Auditory | 0.0767 | 0.6154 | HandSM | 0.0645 | 0.5764 |
| MedPar | 0.0331 | 0.4775 | FaceSM | 0.0368 | 0.6424 |
| ParOcc | 0.0551 | 0.3377 | Auditory | 0.081 | 0.6923 |
| - | - | - | AntMTL | 0.0369 | 0.7892 |
| - | - | - | PostMTL | 0.001 | 0.8162 |
| - | - | - | ParMemory | 0.0425 | 0.4356 |
| - | - | - | Context | 0.0275 | 0.7632 |
| - | - | - | FootSM | 0.0555 | 0.7403 |

**Supplementary Figure 1.**
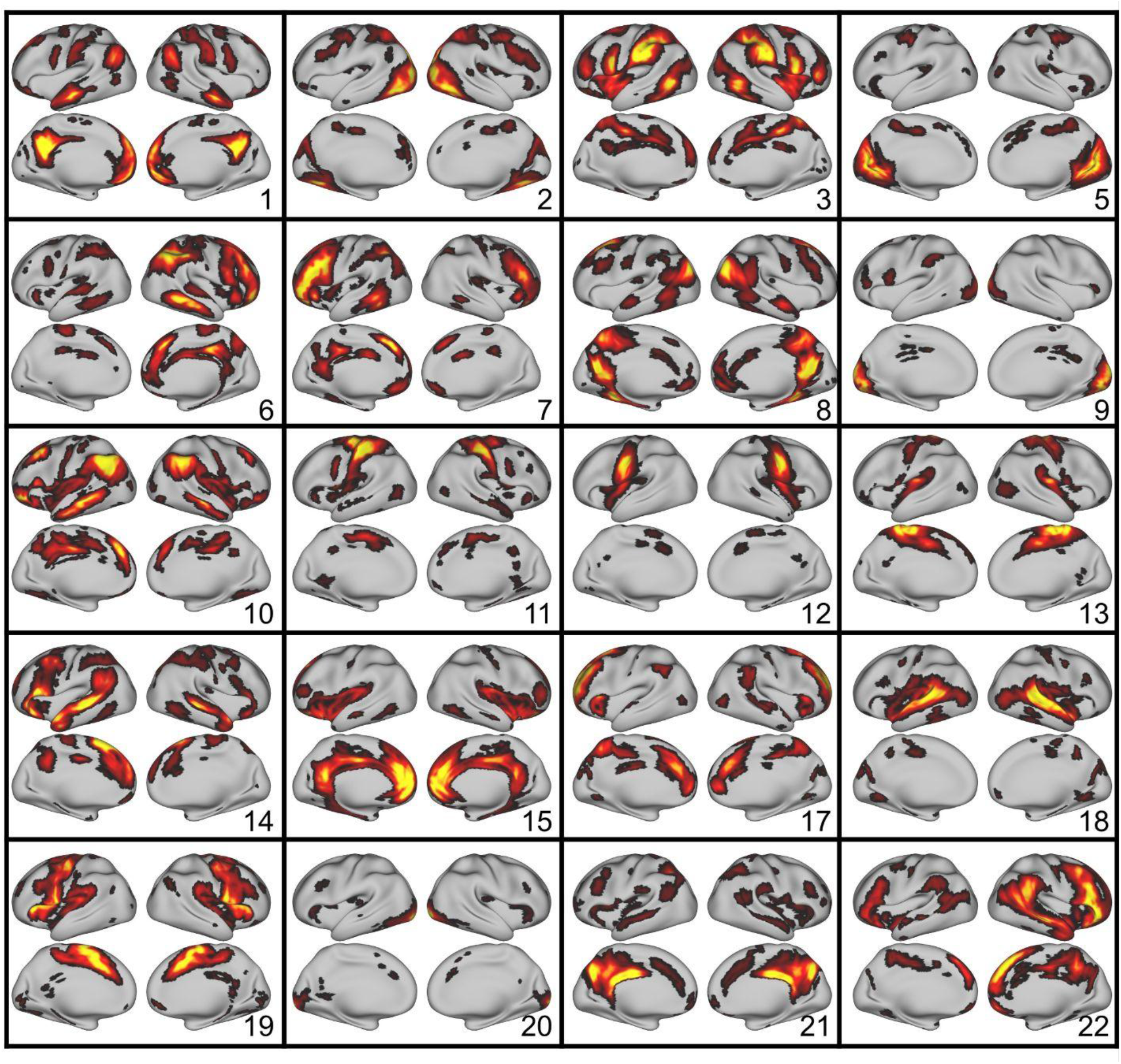
The UKB ICA z-stat maps with 21 good components (https://www.fmrib.ox.ac.uk/ukbiobank/group_means/rfMRI_GoodComponents_d25_v1.txt) were thresholded by FSL melodic mixture-modeling threshold 0.6. The component 16 corresponds to cerebellum and was further excluded, resulting in 20 thresholded UKB ICA maps.

**Supplementary Figure 2.**
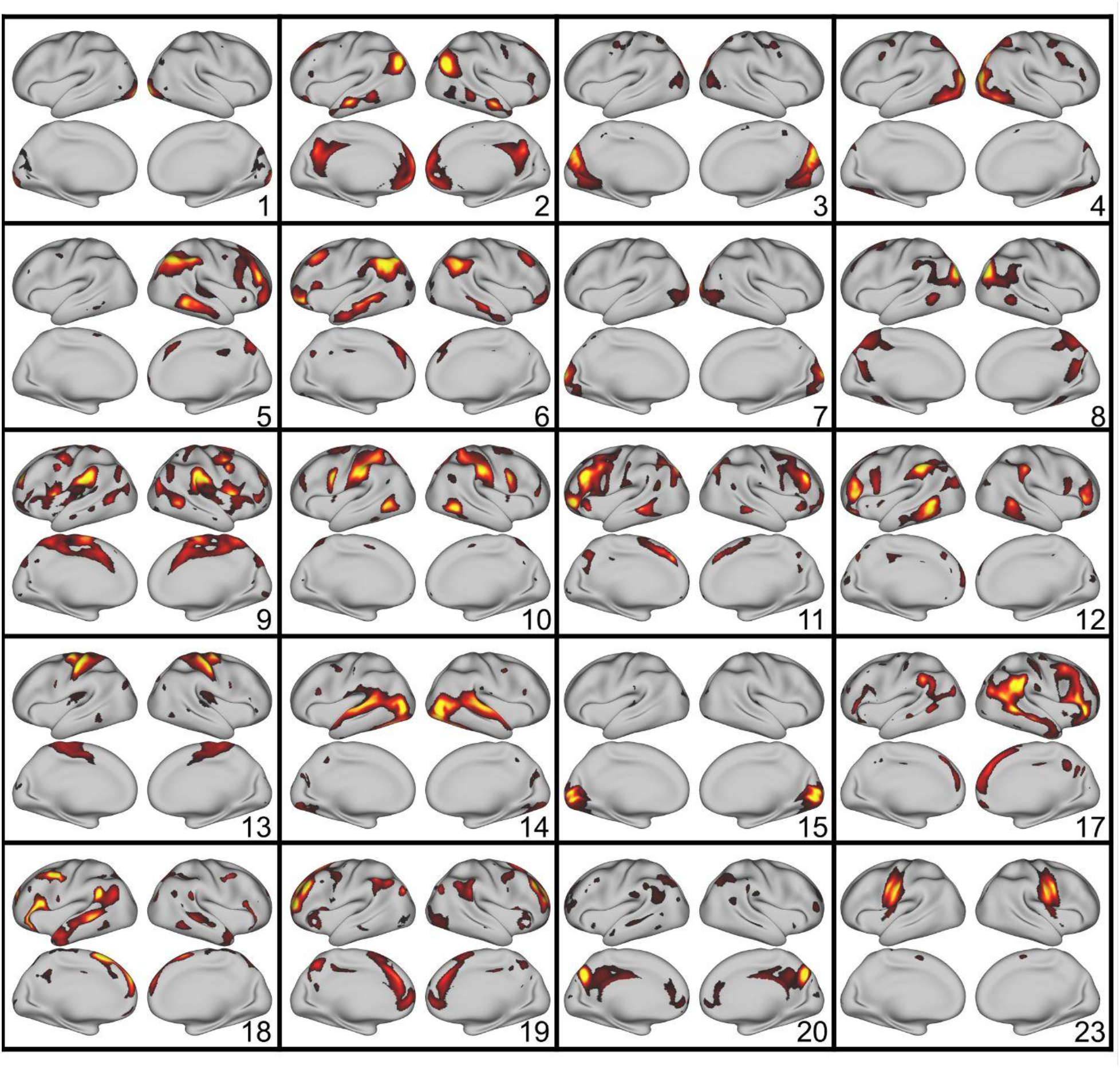
The HCP ICA z-stat maps with 20 cortical components. The HCP group ICA-25 maps “melodic_IC.dscalar.nii” from HCPS1200 release were thresholded by FSL melodic mixture-modeling threshold 0.6. Five components (16, 21, 22, 24, 25) corresponding to subcortical regions were excluded, resulting in 20 thresholded HCP ICA maps.

